# Orphan quality control shapes network dynamics and gene expression

**DOI:** 10.1101/2022.11.06.515368

**Authors:** Kevin G. Mark, SriDurgaDevi Kolla, Danielle M. Garshott, Brenda Martínez-González, Christina Xu, David Akopian, Diane L. Haakonsen, Stephanie K. See, Michael Rapé

## Abstract

All eukaryotes require intricate protein networks to translate developmental signals into accurate cell fate decisions. Mutations that disturb crucial interactions between network components often result in disease, but how the composition and dynamics of complex networks are established is unknown. Here, we identify the tumor suppressor E3 ligase UBR5 as a quality control enzyme that helps degrade unpaired subunits of multiple transcription factors that operate within a single network. By constantly turning over orphan subunits, UBR5 forces cells to continuously replenish network components through new protein synthesis. The resulting cycles of transcription factor synthesis and degradation allow cells to effectively execute the gene expression program, while remaining susceptible to environmental signals. We conclude that orphan quality control plays an essential role in establishing the dynamics of protein networks, which may explain the conserved need for protein degradation in transcription and offers unique opportunities to modulate gene expression in disease.

## Introduction

Metazoan development depends on the formation of protein complexes that differ widely in their stability, stoichiometry, and composition (Huttlin et al., 2021; Li et al., 2021; Padovani et al., 2022). While some interactions between enzymes and substrates only persist for seconds (Pierce et al., 2009), large protein complexes such as the nuclear pore can exist for years (Toyama et al., 2013). When the same complex contains both stable and rapidly exchanging subunits, it is often the transiently bound component that provides important points of regulation (Li et al., 2021; Liu et al., 2018).

Cells use multiple mechanisms to establish interactions at different time scales. Persistent binding is often based on the complementary recognition of amino acid side chains or hydrophobic surfaces of folded domains (Greber and Nogales, 2019; Harper and Schulman, 2021). In contrast, dynamic complexes can rely on intrinsically disordered regions (IDRs) that engage their partners with weak affinity but gain avidity through multivalent binding to several proteins at the same time (Hong et al., 2020; Snead and Gladfelter, 2019). Interactions are further modulated by post-translational modifications, including phosphorylation or ubiquitylation (Ali et al., 2019; Magits and Sablina, 2022), or by small molecules, such as plant hormones or metal ions that can mediate substrate recognition by E3 ligases (Manford et al., 2021; Tan et al., 2007). Developmental signaling requires that distinct types of complex formation are integrated into a coherent response, but how this is accomplished is not well understood.

Accurate complex formation is particularly important for the transcriptional programs that specify cell fate. To read out their target motifs in chromatin, many transcription factors dimerize through Zinc-fingers, BTB domains or leucine zippers (Busch and Sassone-Corsi, 1990; Lourenco et al., 2021). As shown for the pluripotency factors OCT4 and SOX2, dimerization can occur during the search for target motifs, and the complex falls apart when OCT4 and SOX2 dissociate from DNA (Chen et al., 2014). Transcription factors of the BTB family use an alternative strategy and co-translationally form homodimers that are stable for days (Bertolini et al., 2021; Mena et al., 2020). As aberrant heterodimerization impedes the function of BTB proteins, cells have evolved protective pathways to eliminate mispaired regulators of gene expression (Mena et al., 2020; Mena et al., 2018; Padovani et al., 2022). Illustrating the importance of such dimerization quality control, its loss interferes with the development of the peripheral and central nervous systems (Mena et al., 2018).

In addition to their dimerization motifs, transcription factors contain activation domains that are rich in IDRs (Boija et al., 2018; Lu et al., 2018; Trojanowski et al., 2022). A multiplicity of binding sites within their flexible IDRs allows transcription factors to recruit many proteins and thereby nucleate networks that are referred to as transcription, factories, hubs, or condensates (Lourenco et al., 2021; Papantonis and Cook, 2013; Rippe and Papantonis, 2022; Wei et al., 2020). The recruitment of c-MYC, OCT4, or the oncogenic fusion EWS-FLI1 into transcription hubs improves their ability to stimulate gene expression and controls cell fate decisions during development or disease (Boija et al., 2018; Chong et al., 2022; Osborne et al., 2007). Transcription hubs enrich components of the gene expression machinery by up to 1000-fold over nucleoplasmic levels (Jackson et al., 1993), yet they remain dynamic and are rapidly remodeled in response to changing cellular needs (Wei et al., 2020). However, even subtle increases in protein levels can disturb the function of transcription hubs (Chong et al., 2022), and reduced dynamics of networks containing RNA-binding proteins has been associated with disease (Patel et al., 2015; Wagh et al., 2021). How cells establish the proper composition and dynamics of intricate protein networks, such as transcription hubs, is still unknown.

Here, we report our discovery that cells exploit orphan quality control to shape network dynamics and thereby regulate gene expression. Having found that the tumor suppressor UBR5 preserves stem cell pluripotency (Oh et al., 2020), we now show that it acts by helping to degrade unpaired subunits of multiple transcription factors that operate within a single network. By eliminating orphan transcription factors, UBR5 forces cells to constantly replenish network building blocks through new protein synthesis. Although recurrent cycles of protein synthesis, complex formation, and orphan degradation are costly, they allow cells to effectively move through the gene expression program while still being able to shut down transcription in response to stress. We conclude that orphan protein quality control plays a crucial role in establishing dynamic protein networks that can faithfully transmit changes in cell state to the gene expression machinery defining cell fate.

## Results

### UBR5 binds multiple regulators of gene expression

Using human embryonic stem cells (hESC) that express endogenously GFP-tagged OCT4, we recently identified regulators of hESC self-renewal (Oh et al., 2020). Among these, the E3 ligase UBR5 was particularly intriguing: in addition to its function in preserving pluripotency, inactivating mutations in *UBR5* drive mantle cell lymphoma (Meissner et al., 2013), while breast or ovarian cancers amplify the *UBR5* gene to support tumor growth and metastasis (Liao et al., 2017; Qiao et al., 2020; Song et al., 2020). How a single E3 ligase can regulate stem cell identity and act as either a tumor suppressor or oncogene is unknown.

To dissect how UBR5 controls development or disease, we searched for binding partners of endogenous UBR5 that could reveal cellular pathways controlled by this E3 ligase. We used CRISPR-mediated genome engineering to append FLAG epitopes to the amino-terminus of the *UBR5* locus in 293T and HeLa cells, two cell lines that can be grown more easily than hESCs and thus allowed us to isolate endogenous interactors by affinity purification and CompPASS mass spectrometry (Huttlin et al., 2021). Having validated new interactions in transformed cell lines, we could then ask whether UBR5 also engaged these partners in hESCs to identify roles of this E3 ligase at the interface of pluripotency and cancer.

Our purifications of endogenous UBR5 showed that it binds several subunits of the INO80 complex that sustains open chromatin at enhancers of pluripotency factors and oncogenes (Jungblut et al., 2020; Wang et al., 2014; Zhou et al., 2016) (**Figure 1A**). UBR5 also engaged both SPT4 and SPT5 components of the DSIF complex that promotes processive transcript elongation and stem cell differentiation (Aoi et al., 2021; Baluapuri et al., 2019; Hu et al., 2021a; Tastemel et al., 2017). In addition, UBR5 associated with the BUB1, BUBR1, and BUB3 proteins that help establish the mitotic checkpoint complex (MCC), the mitotic kinase PLK1, and the kinesin-13 family member KIF2A. The MCC regulates the anaphase promoting complex (APC/C) that controls nucleosome occupancy at transcription start sites of pluripotency genes (Oh et al., 2020), and overexpression of MCC subunits can drive tumorigenesis (Guardavaccaro et al., 2008; Sotillo et al., 2007; Sotillo et al., 2010).

**Figure 1:**
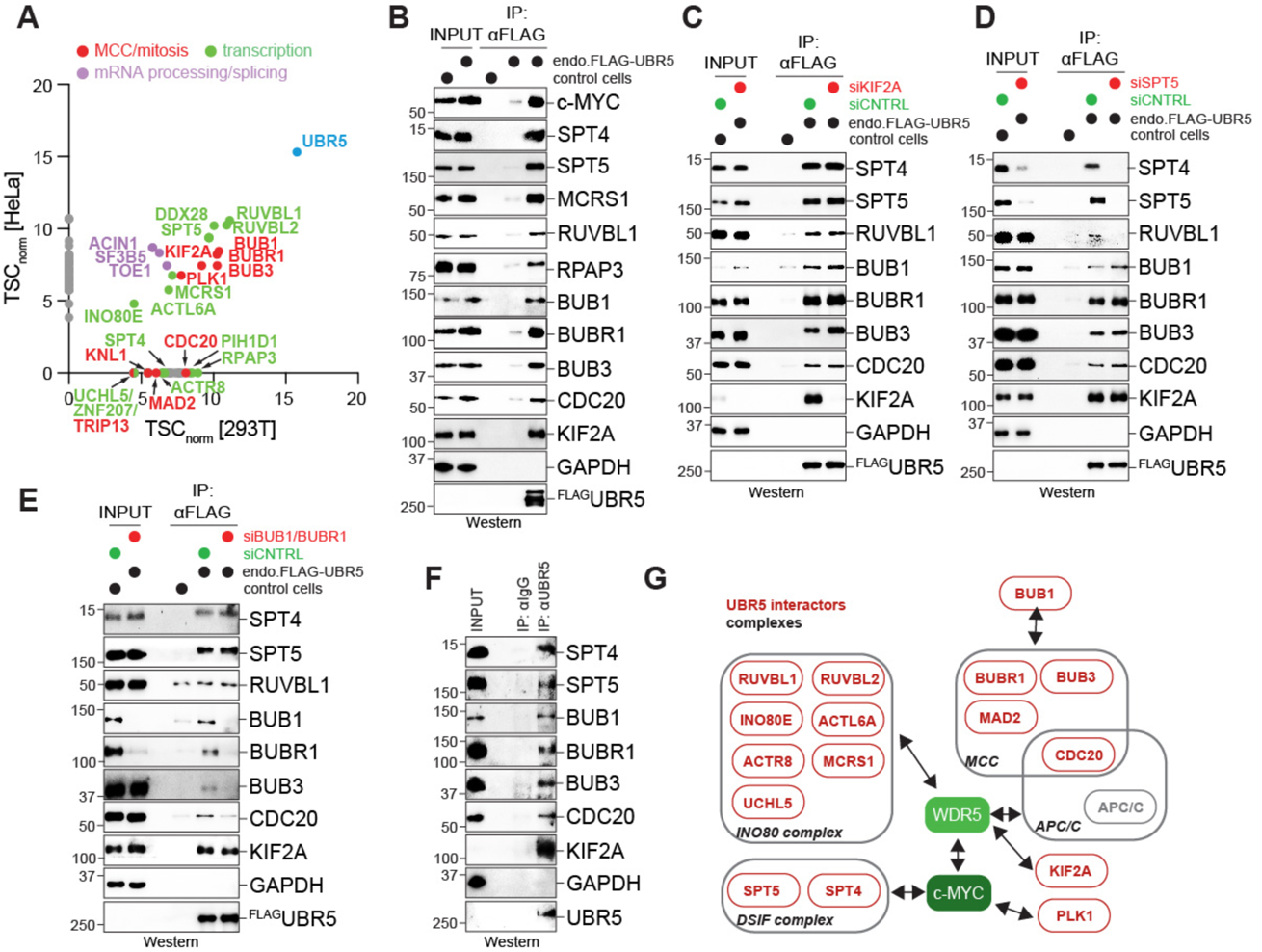
UBR5 binds multiple complexes with tight connections to c-MYC. **A.** Identification of UBR5 binding partners from 293T and HeLa cells expressing endogenously FLAG-tagged UBR5. ^FLAG^UBR5 was affinity-purified from each cell line and binding partners were determined by CompPASS mass spectrometry. Total spectral counts were normalized to UBR5. **B.** Validation of UBR5 binding partners in HeLa cells. Endogenous ^FLAG^UBR5 was immunoprecipitated from HeLa cell lysates and co-purifying proteins were determined by Western blotting using specific antibodies. **C.** Depletion of KIF2A only reduces this protein, but not DSIF or MCC components, in affinity-purifications of endogenous ^FLAG^UBR5 from HeLa cells. **D.** Depletion of SPT5 also reduces the DSIF component SPT4, but not KIF2A or MCC components, in affinity-purifications of endogenous ^FLAG^UBR5 from HeLa cells. **E.** Depletion of BUB1 and BUBR1 only reduces the MCC, but not DSIF or KIF2A, in affinity-purifications of endogenous ^FLAG^UBR5 from HeLa cells. **F.** UBR5 binding partners have close links to the c-MYC/WDR5 complex.

Affinity-purification coupled to Western blotting confirmed the association of endogenous UBR5 with subunits of INO80, DSIF, MCC, and KIF2A (**Figure 1B**). UBR5 forms discrete complexes with these partners: while KIF2A depletion only eliminated this protein from UBR5 immunoprecipitates (**Figure 1C**), loss of SPT5 abrogated binding of the DSIF subunit SPT4 but did not affect recognition of the MCC or KIF2A (**Figure 1D**). Conversely, a reduction in BUB1 and BUBR1 prevented UBR5 from recognizing other MCC components but did not impact binding to DSIF or KIF2A (**Figure 1E**). Importantly, by using antibodies specific for UBR5, we could show that the endogenous E3 ligase engaged the same protein complexes in hESCs (**Figure 1F**).

INO80, APC/C, and KIF2A all engage WDR5 (Ali et al., 2017; Oh et al., 2020; Wang et al., 2014), a methylhistone binding protein that recruits c-MYC to target genes (Thomas et al., 2015). SPT5 and MCRS1 bind c-MYC and N-MYC, respectively (Baluapuri et al., 2019; Jimenez Martin et al., 2021). In addition, PLK1 was reported to affect c-MYC stability and function (Littler et al., 2019), while UBR5 had been proposed to directly target c-MYC for degradation (Qiao et al., 2020; Schukur et al., 2020). Indeed, we find that endogenous UBR5 binds to c-MYC (**Figure 1B**). UBR5 therefore engages many protein complexes with links to c-MYC (**Figure 1G**), a transcription factor that helps establish and maintain stem cell identity and is frequently mutated in cancer (Apostolou and Stadtfeld, 2018; Baluapuri et al., 2020).

### UBR5 degrades multiple transcriptional regulators

UBR5 possesses a UBA domain that binds ubiquitin and a HECT-domain that catalyzes ubiquitin transfer (Kim et al., 2021). These domains allow UBR5 to build K11/K48- or K63/K48-branched ubiquitin chains that elicit efficient proteasomal degradation (Kolla et al., 2022; Ohtake et al., 2018; Yau et al., 2017). To test if UBR5 induces the turnover of proteins that cooperate with c-MYC, we fused each interactor and several of their related proteins to GFP and co-expressed them with mCherry that was under control of an internal ribosome entry site. The ratio between GFP and mCherry, which can be measured by fluorescence associated cell sorting (FACS), reports on the stability of the GFP-tagged protein (Koren et al., 2018; Manford et al., 2020; Sievers et al., 2018). Transcriptional regulators that are turned over through UBR5 should display a higher GFP to mCherry ratio upon depletion of the E3 ligase by siRNAs.

This screen revealed 21 potential targets of UBR5 (**Figure 2A; Figure S1A**). In addition to c-MYC, which had previously been reported to be degraded through UBR5 (Qiao et al., 2020; Schukur et al., 2020), UBR5 depletion stabilized both SPT4 and SPT5 subunits of DSIF, the INO80B, INO80C, INO80F, MCRS1, or RUVBL2 components of INO80, and the MCC subunit CDC20. UBR5 was also needed for the degradation of transcription factors that control cell fate and cooperate with c-MYC, such as OCT4, MAFF, NFIL3, NRL, and TAF1A, or that are activated by stress, including ATF3 and CHOP. In addition, UBR5 helps eliminate the SWI/SNF component SMARCB1, which inhibits c-MYC and whose levels must be calibrated to retain pluripotency (Carmel-Gross et al., 2020; Weissmiller et al., 2019). UBR5 therefore not only binds, but also helps degrade many transcriptional regulators that have links to c-MYC and act at the interface of stem cell and cancer biology.

**Figure 2:**
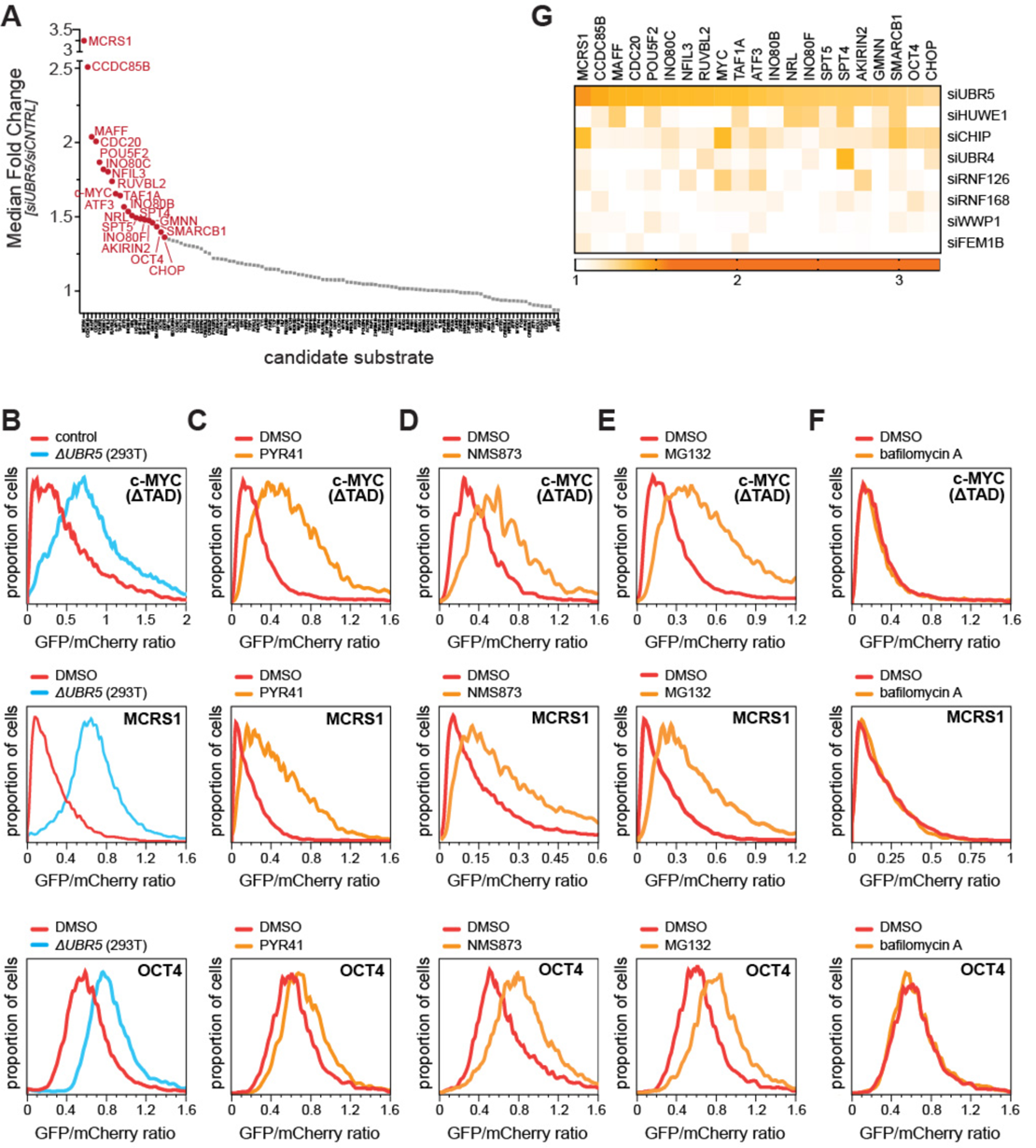
UBR5 degrades multiple transcriptional regulators with tight connections to c-MYC. **A.** Focused screen to identify UBR5 substrates. Subunits of UBR5-interacting complexes as well as their related factors were expressed as GFP-tagged reporters together with mCherry in cells treated with control siRNA or siRNAs targeting UBR5. Protein abundance was measured as the ratio between GFP and mCherry, as determined by FACS. **B.** Validation of select candidate targets (c-MYC^ΔTAD^; MCRS1; OCT4) as GFP-tagged reporters in either control or *ΔUBR5* cells by FACS. A c-MYC reporters lacking the transactivation domain behaved much better in cells, they were used in these experiments. **C.** UBR5 targets are stabilized upon treatment of cells with the E1 enzyme inhibitor PYR41. Reporter levels were determined by FACS. **D.** UBR5 targets are stabilized upon treatment of cells with the p97 inhibitor NMS873. Reporter levels were determined by FACS. **E.** UBR5 targets are stabilized upon treatment of cells with the proteasome inhibitor MG132. Reporter levels were determined by FACS. **F.** UBR5 targets are not affected by lysosomal inhibition through bafilomycin A. Reporter levels were determined by FACS. **G.** Stability of top UBR5 substrates were determined in cell lines that lacked E3 ligases that are functionally or structurally related to UBR5. With few exceptions, notably SPT4, transcriptional regulators are most strongly stabilized by loss of UBR5.

To validate this screen, we inactivated *UBR5* by CRISPR-mediated genome editing in two cell lines and found that candidate substrates were also stabilized by gene deletion (**Figure 2B; Figure S1B**). As expected for a HECT-family E3 ligase, efficient degradation through UBR5 required the E1 enzyme UBA1 and the Cys-specific E2 UBE2L3 (**Figure 2C; Figure S2A, B**). Consistent with UBR5 synthesizing branched ubiquitin chains, inhibition of the main effector of such conjugates, p97 (Yau et al., 2017), stabilized all targets (**Figure 2D; Figure S2C**). By treating cells with MG132 or bafilomycin A, we found that UBR5 substrates were turned over through the proteasome, and not the lysosome (**Figure 2E, F; Figure S2D, E**).

As an additional test for specificity, we measured the stability of each target in cells depleted of E3 ligases that are functionally or structurally related to UBR5. We focused on HUWE1, UBR4, or RNF126 as enzymes that cooperate with UBR5 to eliminate misfolded nascent proteins (Rodrigo-Brenni et al., 2014; Yau et al., 2017); CHIP as an E3 ligase that engages chaperones (Meacham et al., 2001; Scaglione et al., 2011); WWP1 as another HECT E3 ligase with roles in stem cell biology (Hu et al., 2021b); RNF168 as an enzyme identified in the same pluripotency screen as UBR5 (Oh et al., 2020); and FEM1B as an unrelated stress response enzyme (Manford et al., 2020). Apart from SPT4 and SMARCB1, which were efficiently stabilized upon loss of UBR4 or CHIP, each substrate was most strongly protected from degradation by depletion of UBR5 (**Figure 2G**). In addition, we repeated the entire screen in HUWE1-depleted cells and found that this E3 ligase preferentially targeted transcription factors that are not linked to c-MYC (**Figure S2F**). Together, these results document that UBR5 plays an important and specific role in degrading many transcriptional regulators with close links to c-MYC.

### UBR5 ubiquitylates transcription factors

To determine whether UBR5 directly ubiquitylates transcriptional regulators, we reconstituted the activity of this E3 ligase *in vitro*. We purified endogenous UBR5 from HeLa cells and incubated it with the E2 UBE2D3 and candidate substrates produced by *in vitro* transcription and translation. It should be noted that the reticulocyte lysate used to synthesize targets contains E3 ligases that cooperate with UBE2D3 and might support ubiquitin chain initiation, a feature that was critical for the discovery of branched chains (Meyer and Rape, 2014; Wickliffe et al., 2011; Williamson et al., 2009). Importantly, UBR5 robustly ubiquitylated subunits within each complex identified above, including the INO80 components MCRS1, INO80C and RUVBL2, the DSIF subunit SPT5, or the MCC subunit CDC20 (**Figure 3A; Figure S3A**), while SPT4 was modified only poorly (**Figure S3A**). In addition, UBR5 decorated multiple transcription factors, such as c-MYC or NFIL3, with ubiquitin chains. As seen with MCRS1, UBR5 also collaborates with UBE2L3, the E2 enzyme that we found contributes to UBR5-dependent degradation in cells (**Figure 3B; Figure S3B**).

**Figure 3:**
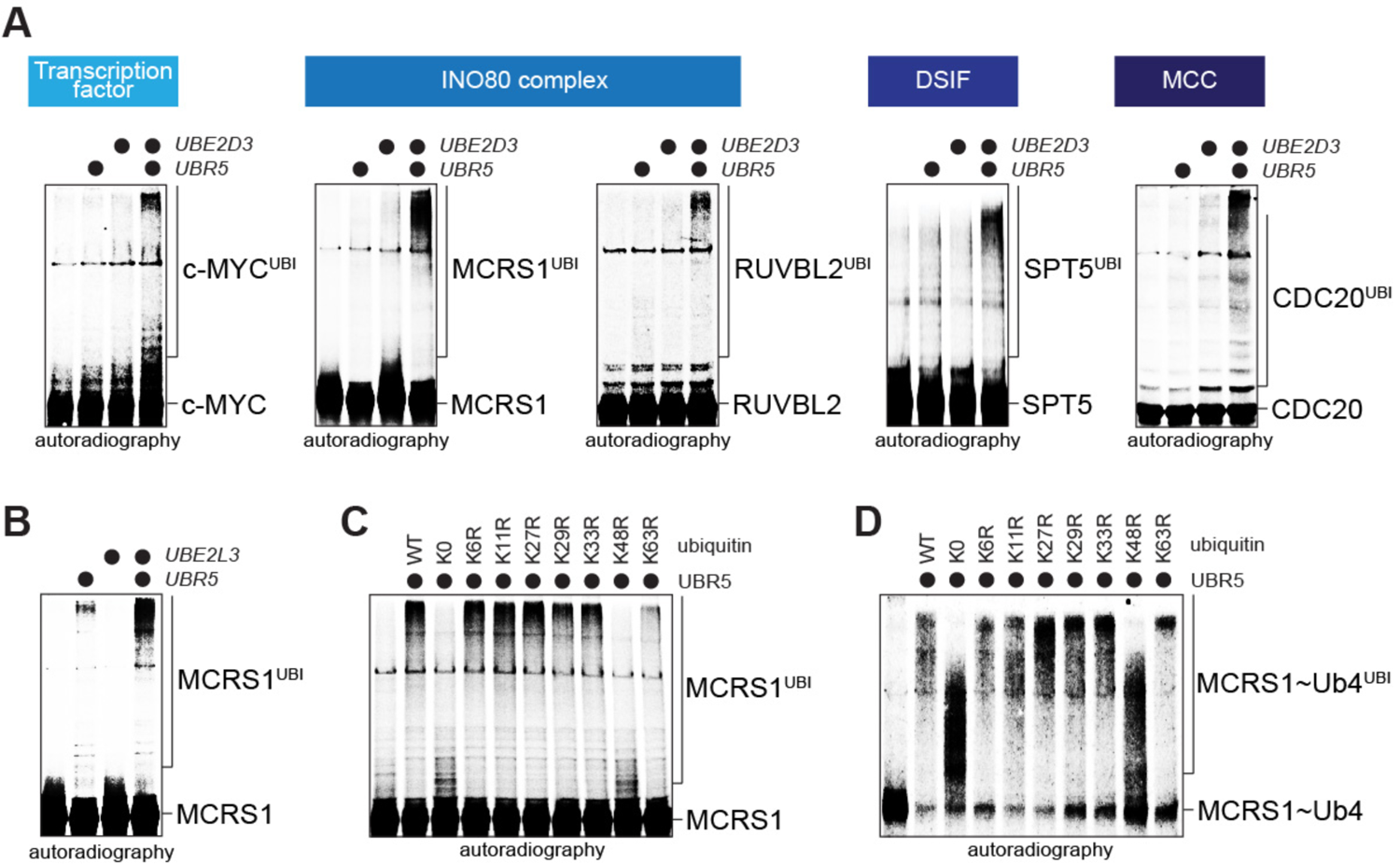
UBR5 modifies multiple transcriptional regulators with proteolytic ubiquitin chains. **A.** UBR5 and the E2 UBE2D3 ubiquitylate transcriptional regulators *in vitro*. Endogenous UBR5 was purified from HeLa cells and incubated with E1, UBE2D3, ubiquitin and ^35^S-labeled targets produced by IVT/T. Substrate modification was analyzed by autoradiography. **B.** UBR5 can cooperate with the E2 UBE2L3. ^35^S-labeled MCRS1 was incubated with UBR5, E1, UBE2L3, and ubiquitin and analyzed for ubiquitylation as above. **C.** UBR5 modifies substrates produced by IVT/T with ubiquitin chains containing K48- and K63-linkages. ^35^S-labeled MCRS1 was incubated with UBR5, E1, UBE2D3, and ubiquitin mutants, and analyzed for ubiquitylation as above. **D.** UBR5 rapidly modifies pre-initiated MCRS1 with K48-linked chains. ^35^S-labeled MCRS1-Ub_4_ was incubated with UBR5, E1, UBE2D3, and ubiquitin mutants and analyzed for ubiquitylation as described above.

Using MCRS1 as our model, we asked whether UBR5 produces ubiquitin chains that can trigger protein degradation through the 26S proteasome. Such conjugates include canonical K48-linked chains or heterotypic K11/K48- and K63/K48-branched conjugates (Kolla et al., 2022; Yau and Rape, 2016). By employing ubiquitin variants that lacked specific Lys residues, we found that UBR5 showed a preference for synthesizing K48-linkages, but chain formation was also reduced if Lys63 of ubiquitin was mutated (**Figure 3C**). Conversely, ubiquitylation was strongly enhanced if we bypassed chain initiation by fusing ubiquitin to MCRS1 (**Figure 3D; Figure S3C**), and pre-initiated MCRS1 was modified by UBR5 in a more K48-specific manner (**Figure 3D**). These results implied that UBR5 branches K48-linked conjugates off chains that contain K63-linkages, and the ubiquitylated MCRS1 was accordingly captured by the effector of branched chains, p97 (**Figure S3D**). We conclude that UBR5 directly acts on transcriptional regulators and modifies them with ubiquitin conjugates that induce very efficient proteasomal degradation.

### UBR5 is required for accurate gene expression

By helping degrade transcriptional regulators, UBR5 might simply limit gene expression, as it had been suggested for select c-MYC targets (Schukur et al., 2020). To test this notion in an unbiased manner, we performed RNAseq in *ΔUBR5* cells and found that a lack of UBR5 indeed upregulated the expression of multiple genes (**Figure 4A**). We confirmed increased expression of select genes by quantitative RT-PCR (**Figure 4B**). Genes that were induced upon *UBR5* deletion included c-MYC targets (**Figure 4A**) and genes with known links to lymphoma (**Figure 4C**), a cancer that results from inactivation of *UBR5* (Meissner et al., 2013).

**Figure 4:**
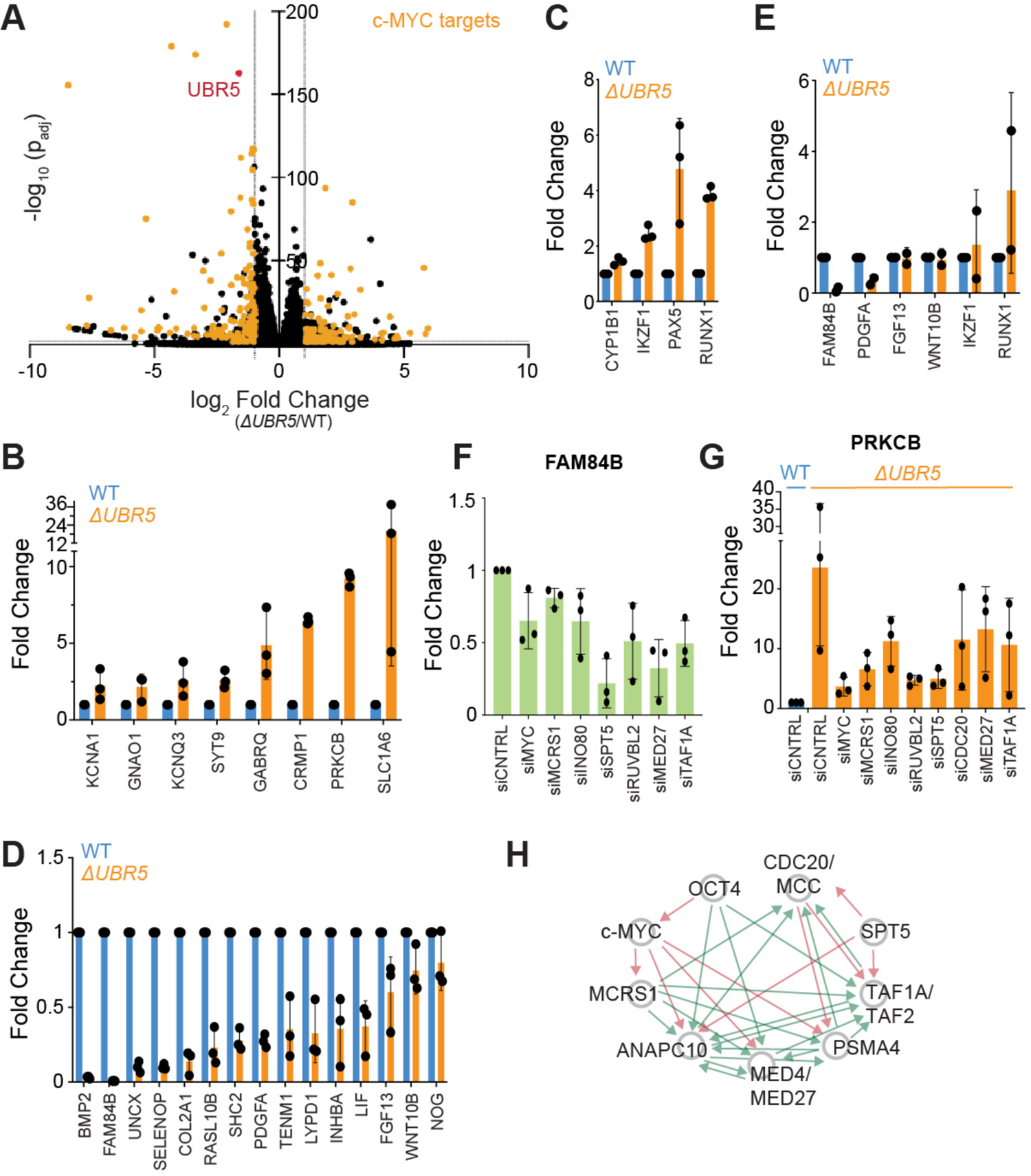
UBR5 is required for accurate gene expression. **A.** RNAseq analysis of control and *ΔUBR5* 293T cells showing that multiple genes are down- and upregulated after *UBR5* deletion. Known target genes of c-MYC are colored in yellow. **B.** Confirmation of select upregulated genes in *ΔUBR5* cells by qRT-PCR. **C.** Upregulation of lymphoma associated genes in *ΔUBR5* cells, as determined by qRT-PCR. **D.** Confirmation of downregulated genes in *ΔUBR5* cells by qRT-PCR. **E.** Similar genes are dependent of UBR5 in hESCs depleted of UBR5. Gene expression was determined after acute UBR5 depletion by siRNAs using qRT-PCR. **F.** Partial depletion of UBR5 substrates and additional network components reduces expression of UBR5-dependent genes, as determined by qRT-PCR. **G.** Partial depletion of UBR5 substrates and additional network components blunts expression of genes that are upregulated upon *UBR5* deletion, as determined by qRT-PCR. **H.** DepMap connections between UBR5 substrates, transcriptional regulators, and additional network components, such as general transcription factors (TAF1A/TAF2), mediator (MED4/MED27), APC/C (ANAPC10), and proteasome (PSMA4). Arrows are shown if correlations are within the top 100 of a respective gene. Negative correlations are shown in red, positive correlations are shown in green.

However, we found that loss of UBR5 also diminished the transcription of a similar number of genes (**Figure 4A**), and many of these UBR5-dependent genes were targets of c-MYC. While our RNAseq analysis was performed in 293T cells, genes that were dependent on UBR5 for full expression included growth factors critical for stem cell pluripotency, such as BMP2, LIF, FGF13, or WNT10B (**Figure 4D**). We found that similar genes were dysregulated in hESCs, although the partial depletion accomplished by siRNAs resulted in less dramatic phenotypes (**Figure 4E**). Thus, UBR5 plays a more nuanced role in gene expression than anticipated: while it restricts the transcription of some genes, it is needed to ensure sustained expression of others.

We next asked if the substrates of UBR5, most of which are known to regulate transcription, modulate the expression of similar genes. We depleted c-MYC, MCRS1, INO80, SPT5, RUVBL2, and CDC20 and measured mRNA levels of UBR5-dependent genes by qRT-PCR. As siRNAs reduce, but do not eliminate their targets, we expected weaker phenotypes than those observed in *ΔUBR5* cells. Still, partial depletion of substrates by siRNAs reduced the expression of UBR5-dependent genes and dampened the effects of *UBR5* loss onto up-regulated genes (**Figure 4F, G; Figure S4A, B**). Consistent with these UBR5 substrates controlling the expression of similar genes, they are highly coordinated with each other across genetic screens cataloged in DepMap and can therefore be considered as components of a functional network (**Figure 4H**). Depletion of additional network components, such as the transcription factor TAF1A or the mediator subunit MED27, also reduced expression of UBR5-dependent genes (**Figure 4G**). We conclude that both transcriptional regulators as well as their degradation through UBR5 are required for accurate gene expression.

### UBR5 targets unpaired c-MYC molecules

Our gene expression analyses indicated that UBR5 is unlikely to simply restrict transcription factor activity. To understand how UBR5 regulates gene expression, we needed to dissect when and how it recognizes its substrates, many of which are subunits of larger complexes. We began this work by identifying the UBR5 degron in c-MYC and then asked whether similar motifs guide the turnover of other components within the UBR5-dependent protein network.

Although previous studies had pointed to an amino-terminal UBR5 degron in c-MYC (Qiao et al., 2020; Schukur et al., 2020), we found that deletion of the entire transactivation domain did not protect c-MYC from UBR5-mediated degradation (**Figure 5A, B**). Further analyses revealed that the carboxy-terminal domain of c-MYC, which contains a helix-loop-helix and a leucine zipper motif, was both necessary and sufficient for turnover by UBR5 (**Figure 5C; Figure S5A**). As UBR5 is a nuclear E3 ligase, the carboxy-terminal domain of c-MYC is only targeted by UBR5 if fused to a nuclear import signal, while a different enzyme mediates its turnover in the cytoplasm (**Figure S5B**).

**Figure 5:**
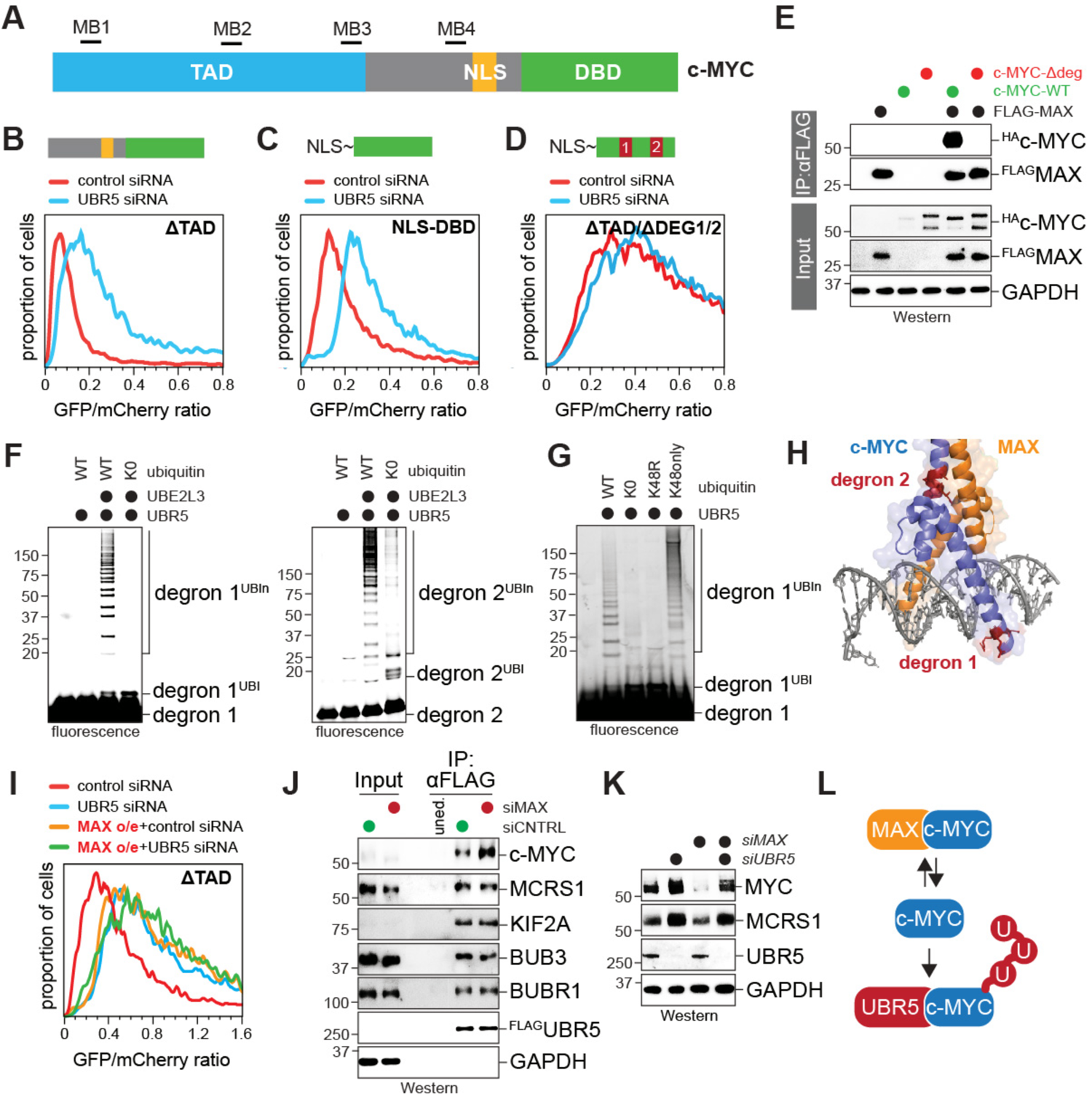
UBR5 targets orphan c-MYC. **A.** Scheme of c-MYC, including the transactivation domain (TAD), the nuclear localization signal (NLS) and the DNA-binding domain (DBD). **B.** UBR5 does not require the TAD of c-MYC to target the transcription factor for degradation. Levels of c-MYC^ΔTAD^-GFP::mCherry reporters were analyzed in cells treated with control or UBR5-siRNAs by FACS. **C.** The carboxy-terminal domain of c-MYC is sufficient to mediate UBR5-dependent degradation. The carboxy-terminal DNA binding domain of c-MYC (DBD) was fused to an N-terminal SV40 NLS and cloned into the degradation reporter, and its levels were determined in control and UBR5 siRNA-treated cells as above. **D.** Mutation of two degrons in the basic helix loop helix and leucine zipper motifs protects c-MYC from UBR5-dependent degradation. Degron mutations were introduced into the c-MYC^ΔTAD^ reporter and its levels were analyzed in control and UBR5 siRNA-treated cells by FACS. **E.** Degron mutations and MAX overexpression stabilize c-MYC. Both degron motifs were mutated in full-length ^HA^c-MYC. Wildtype or degron mutant c-MYC were expressed alone or in the presence of ^FLAG^MAX. Where indicated, ^FLAG^MAX was immune-precipitated and co-purifying c-MYC was detected by Western blotting. **F.** Degron motifs in c-MYC are sufficient for UBR5-dependent ubiquitylation. TAMRA-labeled degrons (degron 1 in BHLH; degron 2 in leucine zipper) were incubated with E1, UBE2L3, UBR5, and ubiquitin, and ubiquitylation was detected on a fluorescence reader after gel electrophoresis. **G.** UBR5 assembles K48-linked chains in a completely reconstituted system. The TAMRA-labeled degron 1 peptide was modified by UBR5 and analyzed as above. As indicated, we used lysine-free ubiquitin (K0); K48R-ubiquitin, or ubiquitin possessing K48 as its only Lys (K48only). **H.** Structural model of the c-MYC/MAX complex, showing that degrons (red) are shielded by DNA-binding and heterodimerization. **I.** MAX overexpression stabilizes c-MYC to the same extent as UBR5 depletion. Cells expressing the c-MYC^ΔTAD^ reporter were treated with control or UBR5 siRNA, and MAX was overexpressed as indicated. Reporter stability was determined by FACS. **J.** Depletion of MAX increases binding of endogenous c-MYC, but not other proteins, to UBR5. Endogenous ^FLAG^UBR5 was immunoprecipitated from 293T cells and co-purifying proteins were detected by Western blotting. As indicated, MAX was depleted by siRNAs. Cells were treated with proteasome inhibitors to prevent c-MYC degradation in the absence of MAX. **K.** UBR5 targets orphan c-MYC. Cells were treated with control, MAX, or UBR5 siRNAs and levels of c-MYC were determined by Western blotting. **L.** Model of orphan c-MYC degradation by UBR5.

Based on these results, we performed Ala- and Glu-scans of the carboxy-terminal domain and found that degradation through UBR5 was lost if we simultaneously mutated motifs in the helix-loop-helix region and leucine zipper (**Figure 5D; Figure S5C**). While loss of single motifs had little impact on c-MYC turnover, combination of the respective mutations strongly stabilized c-MYC and protected it from UBR5. The same mutations increased c-MYC levels as detected by Western blotting (**Figure 5E**). Both motifs are conserved in N-MYC, but less so in L-MYC (**Figure S5D**), which is consistent with only the former being targeted by UBR5 (**Figure S5E**).

To test if UBR5 directly recognizes the candidate c-MYC degrons, we synthesized each motif as a TAMRA-labeled peptide and incubated them with UBR5 and UBE2L3. As UBE2L3 can transfer ubiquitin only to the catalytic Cys of UBR5, but not to Lys residues (Wenzel et al., 2011), any degron ubiquitylation observed in this reconstituted system will be specific for UBR5. Importantly, each degron peptide was efficiently modified by UBR5 and UBE2L3 (**Figure 5F**), and UBR5 required K48 of ubiquitin for chain formation (**Figure 5G**). We conclude that c-MYC contains two carboxy-terminal degrons that are each sufficient to mediate recognition by UBR5.

When mapping these motifs onto c-MYC structures, we noted that both degrons become buried when c-MYC forms a functional DNA-bound transcription factor with MAX (**Figure 5H**). In addition, mutation of its degrons disrupted the interaction of c-MYC with MAX (**Figure 5E**), which implied that degron residues in c-MYC can bind MAX or UBR5, but not both proteins at the same time. Supporting this notion, overexpression of MAX prevented the UBR5-dependent degradation of c-MYC, as seen by FACS and Western blot analysis (**Figure 5E, I**). A cancer-relevant mutant of MAX that is impaired in complex formation with c-MYC (Jimenez Martin et al., 2021) stabilized c-MYC less efficiently than the wildtype protein (**Figure S5F**). Depletion of MAX had the opposite effect and stimulated capture of c-MYC by UBR5 (**Figure 5J**), and accordingly decreased c-MYC levels in a UBR5-dependent manner (**Figure 5K**). Because UBR5 requires p97 for substrate degradation, we asked whether depleting the p97 co-adaptor UFD1/NPL4 rescued c-MYC levels in the absence of MAX and found this to be the case as well (**Figure S5G**). In contrast to c-MYC, MAX is not targeted by UBR5 and hence solely acts as a stabilizing partner (**Figure S5H**). Akin to an orphan quality control E3 ligase, UBR5 therefore selectively targets c-MYC molecules that are not bound to MAX (**Figure 5L**).

### Complex formation stabilizes multiple UBR5 substrates

By titrating a c-MYC degron into *in vitro* ubiquitylation reactions, we noted that the modification of MCRS1 and SPT5 by UBR5 was inhibited in a dose-dependent manner (**Figure 6A**). UBR5 therefore engages c-MYC and other network components through an overlapping site, raising the possibility that it acts as an orphan E3 ligase towards additional transcriptional regulators. To test this hypothesis, we expressed UBR5-targets with dimerization partners and monitored effects on UBR5-dependent degradation. Strikingly, we found that complex formation stabilized every protein we tested: MAFF and CDC20 were protected by BACH2 and MCC to the same extent as by UBR5-depletion, and loss of UBR5 did not induce further stabilization when the partner was co-expressed (**Figure 6B; Figure S6A**); such targets mirror c-MYC, as UBR5 is their major orphan E3 ligase. MCRS1, RUVBL2, or OCT4 were strongly stabilized by either co-expression of their partners or UBR5 depletion, yet partners stabilized the target to a stronger extent than loss of UBR5 (**Figure 6C-E)**; these proteins are controlled as orphan subunits by UBR5 as well as other E3 ligases. By contrast, SPT4 was much more stabilized by co-expression of SPT5 than by loss of UBR5 and hence mostly relies on other E3 ligases for orphan quality control (**Figure S6B**). Like MAX, the stabilizing partners SOX2, RUVBL1, or BACH2 were not targeted for degradation by UBR5, which suggests that protective effects were not simply due to competitive inhibition of UBR5 (**Figure S6C**).

**Figure 6:**
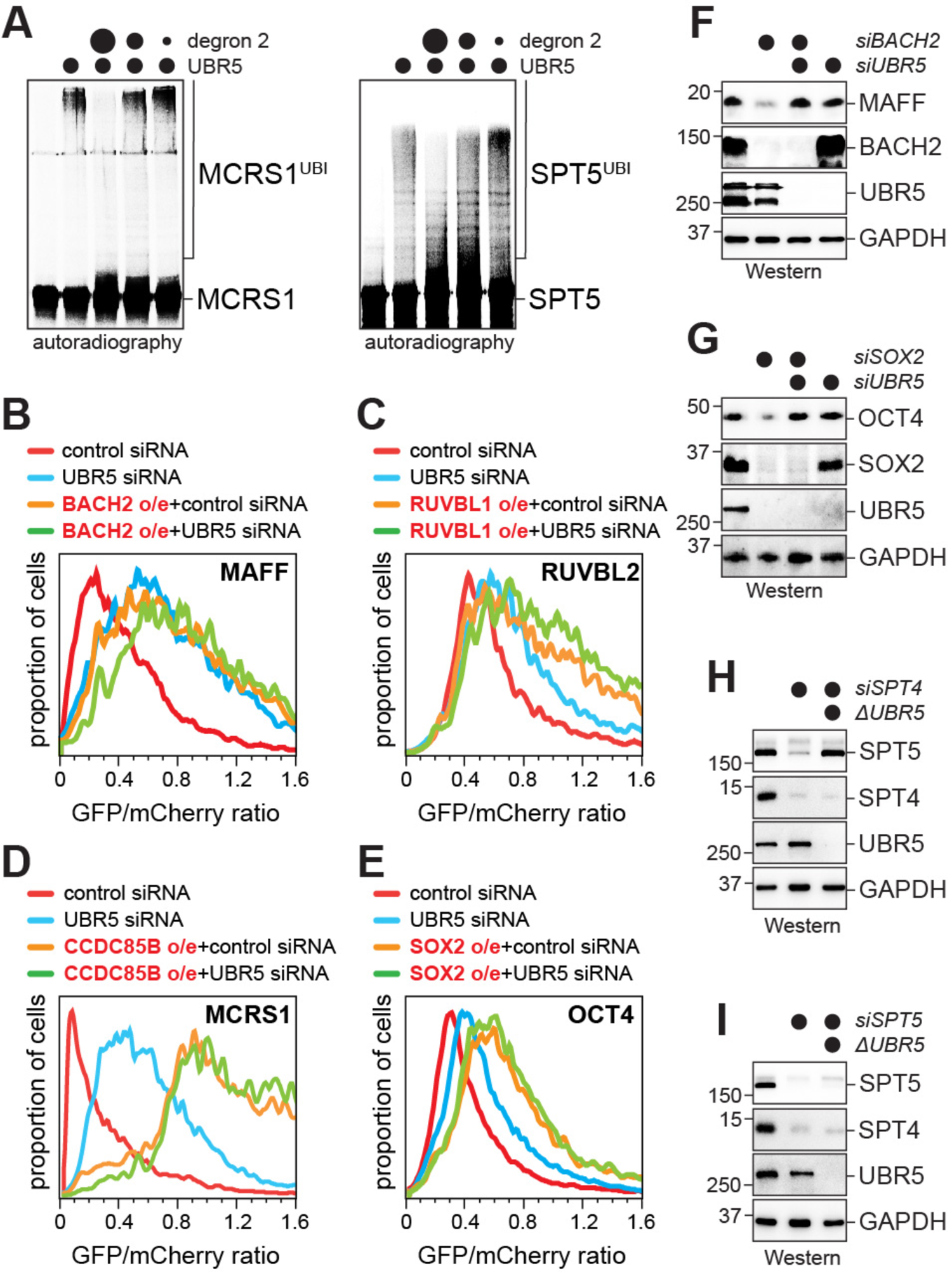
UBR5 targets multiple transcriptional regulators as orphan proteins. **A.** UBR5 binds other transcriptional regulators through the same site as c-MYC. ^35^S-labeled MCRS1 or SPT5 were incubated with E1, UBE2L3, UBR5, ubiquitin, and increasing concentrations of the recombinant c-MYC degron-2 peptide, as indicated. Ubiquitylation was monitored by autoradio-graphy. **B.** BACH2 protects MAFF from UBR5-dependent degradation. The MAFF-reporter was expressed in control cells or cells treated with siRNAs targeting UBR5. As indicated, BACH2 was co-expressed, and the stability of the MAFF reporter was determined by FACS. **C.** The RUVBL2 reporter was stabilized by RUVBL1 co-expression. Reporter stability was assessed as described above. **D.** The MCRS1 reporter was stabilized by CCDC85B expression. Reporter stability was assessed as described above. **E.** The OCT4 reporter was stabilized by SOX2 expression. Reporter stability was assessed as described above. **F.** BACH2 protects endogenous MAFF from UBR5-dependent degradation. 293T cells were depleted of BACH2 and/or UBR5 by using siRNAs, as indicated, and endogenous MAFF were visualized by Western blotting. **G.** SOX2 protects endogenous OCT4 from UBR5-dependent degradation in stem cells. H1 hESCs were depleted of SOX2 and/or UBR5 using siRNAs and levels of OCT4 were determined by Western blotting. **H.** Deletion of *UBR5* stabilizes orphan SPT5. Wildtype or *ΔUBR5* 293T cells were depleted of SPT4, as indicated, and levels of SPT5 were determined by Western blotting. **I.** Deletion of *UBR5* is not sufficient to stabilize orphan SPT4. Wildtype or *ΔUBR5* 293T cells were depleted of SPT5, as indicated, and levels of SPT4 were determined by Western blotting.

While experiments with endogenous proteins were limited by the availability of antibodies, they mirrored our reporter studies. As we had seen with c-MYC, MAFF levels strongly decreased in transformed cell lines, when its partner BACH2 had been depleted, but UBR5 co-depletion by siRNAs restored MAFF (**Figure 6F**). This behavior is conserved across cell types, as OCT4 levels dropped in untransformed hESCs in a UBR5-dependent manner if SOX2 had been depleted (**Figure 6G**). The impact of SOX2 depletion on OCT4 levels was likely tempered by a concurrent loss of UBR5, pointing towards potential feedback regulation that will be explored elsewhere. Substrate behavior was also independent of how we induced UBR5 loss: similar to depletion by siRNAs, *UBR5* deletion by genome editing rescued the drop in SPT5 seen upon loss of its partner SPT4 (**Figure 6H**). In line with our findings that other E3 ligases target SPT4 (**Figure S6B**), *UBR5* deletion did not stabilize SPT4 in the absence of SPT5 (**Figure 6I**). We conclude that UBR5 is an orphan E3 ligase for multiple transcription factors that operate within the same network.

### Orphan transcription factor degradation establishes network dynamics and function

Orphan proteins often arise due to an imbalance in the synthesis of complex subunits or as a result of stresses that alter the efficiency and kinetics of complex formation (Juszkiewicz and Hegde, 2018). If UBR5 only degrades excessive or defective subunits, its loss should increase its targets, while complexes containing functional proteins might not accumulate. If, however, UBR5 plays an additional regulatory role, it might also control the abundance or stability of transcription factor complexes.

To monitor protein or complex stability, we treated cells with the protein synthesis inhibitor cycloheximide. As expected, *UBR5* deletion increased the levels of endogenous c-MYC, but not MAX, and c-MYC was detected for longer times after cycloheximide treatment (**Figure 7A**). Using ^FLAG^MAX as bait, we found that the loss of *UBR5* also strongly increased the abundance and persistence of c-MYC/MAX complexes. A similar rise in complex levels was seen when we purified endogenous MAX from *ΔUBR5* cells (**Figure 7B**). The accumulation of c-MYC and c-MYC/MAX complexes in *ΔUBR5* cells was even more drastic if c-MYC was transiently overexpressed, as expected for a condition that produces more orphan transcription factor subunits (**Figure S7A**). UBR5 therefore restricts the levels of its target, c-MYC, and the major complex containing this protein, c-MYC/MAX.

**Figure 7:**
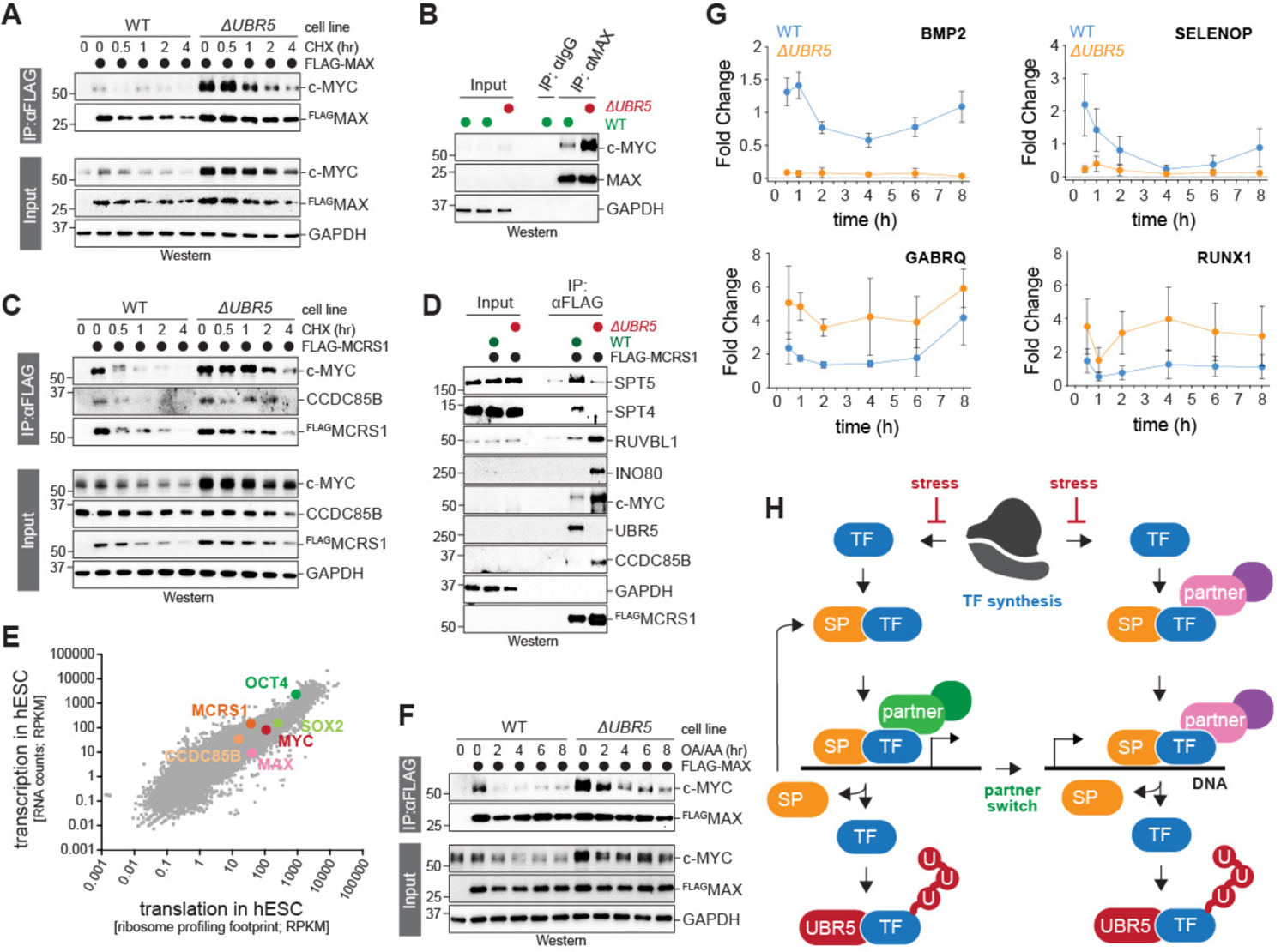
Orphan transcription factor degradation establishes network dynamics and function. **A.** *UBR5* deletion stabilizes c-MYC as well as c-MYC/MAX complexes. Wildtype or *ΔUBR5* cells expressing ^FLAG^MAX were treated with cycloheximide. ^FLAG^MAX was immuno-precipitated and co-purifying proteins were detected by Western blotting using specific antibodies. **B.** *UBR5* deletion stabilizes endogenous c-MYC/MAX complexes. Endogenous MAX was precipitated from wildtype or *ΔUBR5* cells and co-purifying c-MYC was detected by Western blotting. **C.** *UBR5* deletion stabilizes CCDC85B/MCRS1 complexes. ^FLAG^MCRS1 was precipitated from wildtype or *ΔUBR5* cells and co-purifying proteins were detected by Western blotting. Cycloheximide was added as indicated. **D.** *UBR5* deletion prevents MCRS1 from binding to DSIF. ^FLAG^MCRS1 was immunoprecipitated from wildtype or *ΔUBR5* cells and co-purifying proteins were detected by Western blotting. **E.** hESCs produce the degraded subunit in excess over its stabilizing partner. Using a previously published dataset (Werner et al., 2015), transcription was analyzed in H1 hESCs by RNAseq, while translation was followed by ribosome profiling. Unstable subunits are shown in light, stabilizing partners in dark colors. **F.** Mitochondrial stress dismantles c-MYC/MAX complexes. ^FLAG^MAX was immunoprecipitated from wildtype or *ΔUBR5* cells treated with oligomycin and antimycin A, and co-purifying proteins were determined by Western blotting. **G.** Mitochondrial stress decreases expression of UBR5-dependent genes, while independent genes are less affected. Gene expression was analyzed in wildtype or *ΔUBR5* cells after oligomycin and antimycin A treatment using qRT-PCR. **H.** Orphan quality control establishes network dynamics and function. Transcription factors (TF) in the c-MYC network are continuously produced, form complexes with stabilizing partners (SP) and function in gene expression. Following their dissociation, UBR5 degrades one subunit to render complex disassembly irreversible. The stabilizing partner can be re-used in another transcription factor cycle, while the unstable subunit needs to be re-synthesized. This network architecture allows transcription factors to efficiently switch from early to late partners in the gene expression program, while remaining susceptible to stress-induced inhibition of gene expression.

To determine if similar regulation occurs for other network components, we monitored the abundance and interactions of MCRS1, which is required for the expression of UBR5-dependent genes and was our model substrate in biochemical analyses. Deletion of *UBR5* slightly increased MCRS1, but complexes between MCRS1 and its stabilizer CCDC85B persisted much longer in *ΔUBR5* cells (**Figure 7C**). Similar complex stabilization was observed upon reciprocal purification of CCDC85B (**Figure S7B**). Intriguingly, *UBR5* deletion caused c-MYC to accumulate in MCRS1 immunoprecipitates, suggesting that UBR5 also restricts crosstalk between complexes within the larger network. Reduced complex turnover in *ΔUBR5* cells impacted which partners MCRS1 can engage: while in control cells MCRS1 readily co-precipitated with the DSIF complex that acts late in transcript elongation, it failed to bind DSIF in cells lacking *UBR5* (**Figure 7D**). By contrast, MCRS1 was still able to interact in *ΔUBR5* cells with INO80, which acts early in transcription by modulating chromatin architecture. This behavior mirrors c-MYC, which fails to engage regulators of transcription elongation if its ubiquitylation is impaired (Endres et al., 2021), and suggesst that orphan subunit degradation establishes network dynamics that enable transcription regulators to switch from early to late partners.

These experiments also revealed that cells rely on protein synthesis to counteract UBR5 and sustain transcriptional regulators at their functional levels. Underscoring the broad nature of this regulatory feature, we found by RNAseq and ribosome profiling that hESCs produce c-MYC in ∼10-fold excess over MAX (**Figure 7E**), and similar observations were made for MCRS1 and CCDC85B, OCT4 and SOX2, SPT5 and SPT4, or MAFF and BACH2 (**Figure 7E; Figure S7C**). As protein synthesis can be shut off in response to stress (Costa-Mattioli and Walter, 2020), this signaling architecture could couple gene expression to developmental or environmental signaling. Indeed, mitochondrial stress, which inhibits protein synthesis (Guo et al., 2020), depleted c-MYC/MAX complexes, and this response was delayed if *UBR5* had been deleted (**Figure 7F**). The same condition impaired expression of genes that were dependent on UBR5 (*BMP2, SELENOP*), while it had less of an impact on genes that were induced by the absence of the orphan E3 ligase (*GABRQ*, *RUNX1*) (**Figure 7G**). By establishing a dynamic network, orphan quality control therefore also allows the gene expression machinery to remain susceptible to environmental inputs.

## Discussion

Quality control pathways are often thought to eliminate defective proteins to prevent aggregation and tissue degeneration. As the propensity for protein misfolding increases with age, phenotypes of insufficient quality control are mainly observed in older organisms after their reproductive phase (Brehme et al., 2014; Harper and Bennett, 2016; Hipp et al., 2014; Kaushik and Cuervo, 2015; Vilchez et al., 2014), and why quality control systems have been conserved so well through evolution is not fully understood. Here, we show that stem cells use orphan quality control to establish a dynamic protein network that regulates gene expression and cell identity. Loss of the key enzyme, the E3 ligase UBR5, stabilizes interactions within the transcription factor network and compromises the efficiency and adaptability of gene expression. Our findings reveal an essential regulatory role of quality control beyond the removal of toxic protein species, which may explain the conserved need for protein degradation in gene expression and offers exciting possibilities to modulate the transcription machinery for therapeutic benefit.

### UBR5 exerts orphan quality control

We had previously identified UBR5 as an E3 ligase that is required for hESC self-renewal (Oh et al., 2020). Befitting a role in pluripotency, UBR5 is essential for embryogenesis (Saunders et al., 2004) and frequently dysregulated in cancer (Liao et al., 2017; Meissner et al., 2013; Qiao et al., 2020; Schukur et al., 2020; Song et al., 2020), but how UBR5 acts at the interface of development and disease was unknown. Using a substrate discovery approach that integrates mapping of endogenous UBR5 interactors with focused stability measurements, we found that UBR5 controls multiple transcription factors that act within a single gene expression network. Rather than curbing transcription factor activity, UBR5 supports their function by establishing dynamic interactions that are needed for accurate gene expression.

The central transcription factor within the UBR5 network is c-MYC, whose stabilization or overexpression causes cancer (Welcker and Clurman, 2008). Organisms therefore restrict c-MYC accumulation, and previous work had identified several E3 ligases that engage the transactivation domain of c-MYC to limit oncogenic gene expression (Adhikary et al., 2005; Huber et al., 2016; King et al., 2016; Reavie et al., 2010; Welcker and Clurman, 2008; Welcker et al., 2004; Welcker et al., 2022). UBR5 acts differently from these enzymes: it targets degrons in the carboxy-terminal domain of c-MYC that become inaccessible upon formation of DNA-bound c-MYC/MAX dimers. Rather than eliminating a chromatin-bound transcription factor, UBR5 degrades orphan c-MYC molecules that do not engage DNA (**Figure 5L**). In addition to the experiments presented here, c-MYC levels drop upon deletion of *MAX* or following chemical inhibition of c-MYC/MAX complex formation (Blackwood et al., 1992; Hishida et al., 2011; Yao et al., 2022), and we expect that UBR5 degrades c-MYC under these conditions as well. For complex subunits that are unstable as orphan proteins, such as c-MYC, disrupting complex formation to elicit orphan quality control will likely be an effective strategy to accomplish small-molecule induced protein degradation. In addition, compounds that accelerate recognition of orphan proteins by their E3 ligases should increase transcription factor turnover and thereby disrupt pathological gene expression.

UBR5 not only targets c-MYC, but also other transcriptional regulators within the same network, as orphan proteins. Subunits of INO80, DSIF, or the MCC were all protected from UBR5 by co-expression of their partners, and their levels dropped in a UBR5-dependent manner when their interactors were depleted. Competition experiments showed that UBR5 detects these targets through the same site as c-MYC. A parallel study noted that UBR5 binds to steroid receptors through a motif that is also engaged by co-activators (N. Thomä, B. Ebert, submitted), indicating that UBR5 might similarly target other transcription factors once they have released their critical partners. In addition, we previously found that UBR5 ubiquitylates misfolded nascent proteins to prevent their aggregation (Yau et al., 2017). As misfolding impedes complex formation, we anticipate that UBR5 also functions as an orphan E3 ligase in this context. We therefore propose that UBR5 is a canonical orphan quality control E3 ligase.

Given the substrate specificity of UBR5 and its dysregulation in cancer, our work provides a firm link between quality control and tumorigenesis. *UBR5* deletion increases the expression of lymphoma-associated genes, which might contribute to its role as a tumor suppressor in mantle cell lymphoma (Meissner et al., 2013). By contrast, breast and ovarian cancers amplify *UBR5* (Qiao et al., 2020; Schukur et al., 2020; Song et al., 2020). These tumors are characterized by a high degree of aneuploidy, which disrupts coordinated expression of subunits encoded on distinct chromosomes and leads to a broad rise in orphan proteins (Brennan et al., 2019; Siegel and Amon, 2012; Torres et al., 2007). As untransformed hESCs already produce transcription factor subunits in strong excess over their stabilizing partners (**Figure 7E**), a further imbalance in the synthesis of complex subunits through aneuploidy could result in orphan protein levels that activate stress pathways preventing cancer initiation or survival (Bartkova et al., 2005). By eliminating orphan proteins, UBR5 therefore not only ensures accurate gene expression but may also protect cancer cells from aneuploidy-induced proteotoxic stress. We propose that small molecules that inhibit UBR5 and thus curb orphan quality control should be tested for therapeutic benefit in models of aneuploid breast or ovarian cancer with *UBR5* amplification.

### Orphan quality control promotes gene expression

As orphan proteins often expose hydrophobic surfaces (Padovani et al., 2022; Zavodszky et al., 2021), we had initially hypothesized that orphan transcription factor degradation protects the gene expression machinery from being captured in non-productive aggregates. However, we could not detect aggregates of c-MYC or other targets, even if these proteins were overexpressed in *ΔUBR5* cells. Moreover, we found that c-MYC and MCRS1 continue to engage their functional partners in *ΔUBR5* cells, which is inconsistent with being diverted to non-specific aggregates.

Instead, our mechanistic work revealed an essential regulatory function for orphan quality control that complements its role in degrading excess proteins. Previous work had revealed many complexes that intersect with c-MYC, such as OCT4/SOX2, INO80, MCC, or DSIF, and we found that UBR5 targets orphan subunits of all these complexes for degradation (Baluapuri et al., 2019; Endres et al., 2021; Guarnaccia and Tansey, 2018; Jimenez Martin et al., 2021; Lourenco et al., 2021; Oh et al., 2020; Thomas et al., 2015; Wang et al., 2014; Zhou et al., 2016). Inactivating these transcriptional regulators phenocopied the loss of c-MYC in gene expression and across many genetic screens, suggesting that they are components of a single network. Our interaction studies showed that the building blocks of this network are continuously dismantled, and orphan subunit degradation by UBR5 renders complex disassembly irreversible and imposes a need to re-synthesize transcription factors. Orphan quality control therefore plays a critical role in creating recurrent cycles of protein synthesis and transcription factor degradation that shape the dynamics of a protein network required for c-MYC dependent gene expression (**Figure 7H**).

Why do cells use such a costly mode of regulation? c-MYC acts at multiple steps in gene expression and engages chromatin modifiers, RNA polymerases, and regulators of transcription elongation (Baluapuri et al., 2020; Lourenco et al., 2021). For such multitasking proteins, the same molecule must either change its interactions as gene expression proceeds from one step to the next or distinct molecules act at subsequent steps of this program. As transcription occurs in hubs of high protein concentrations (Rippe and Papantonis, 2022; Wei et al., 2020), it could be difficult for a molecule to move between binding partners in a coordinated fashion. By contrast, cycles or protein synthesis and degradation allow cells to turn over the first complex and then form the next outside of the hub after the unstable subunit has been re-synthesized. Consistent with this model, abolishing orphan subunit degradation did not prevent c-MYC and MCRS1 from engaging early regulators of transcription, such as INO80, but impaired recognition of the DSIF complex that acts late during transcript elongation (**Figure 7D**) (Endres et al., 2021; Jaenicke et al., 2016). We suggest that orphan quality control allows the gene expression machinery to move through subsequent steps of its complex program despite the high concentrations in transcription hubs.

By imposing a need for protein synthesis to sustain transcription factor complexes, orphan quality control also introduces means for transcription regulation. Protein synthesis is under tight control of the integrated stress response, which prevents cells from producing proteins under adverse conditions (Costa-Mattioli and Walter, 2020). We found that mitochondrial dysfunction, a stress that activates this stress response (Guo et al., 2020), depletes MYC/MAX complexes and impairs expression of genes under control of UBR5. It is likely that other stressors, such as viral infection or proteotoxic overload, similarly interfere with c-MYC dependent gene expression to delay cell division under challenging conditions. By optimizing network dynamics, orphan quality control therefore ensures that the transcription machinery remains sensitive to changes in the cellular environment. We conclude that cells invest in recurrent cycles of transcription factor synthesis and degradation to increase both the efficiency and adaptability of gene expression programs that must integrate environmental or developmental inputs.

The role of protein degradation in establishing network dynamics could explain apparently paradoxical findings that proteolytic enzymes, such as the segregase p97 or the 26S proteasome, are required for transcription (Ferdous et al., 2001; Heidelberger et al., 2018; Huang et al., 2006; Kinyamu and Archer, 2007). In fact, the better a transcription factor stimulates gene expression, the faster it is degraded (Salghetti et al., 2001; Salghetti et al., 2000; Wang et al., 2017; Wu et al., 2007). We propose that rapid degradation optimizes the dynamics of gene expression networks that must integrate environmental or developmental inputs into a coherent output. Our work also sheds light on the observation that nuclear quality control occurs in the nucleolus (Frottin et al., 2019; Fu et al., 2021), which is the site of c-MYC dependent production of ribosomal RNA. We suggest that the nucleolus may already be enriched in orphan E3 ligases that are required for gene expression and could readily be repurposed to eliminate misfolded proteins. By identifying orphan E3 ligases at the intersection of transcription regulation and aggregate protection, such as UBR5, we stand to find many new opportunities to modulate gene expression programs and thereby provide therapeutic relief in diseases caused by transcription factor dysregulation.

## Author contributions

K.G.M., S.K., D.M.G., B.M. and C.X. performed the experiments. D.A. and D.L.H. made CRISPR-edited cell lines. S.K.S. generated recombinant p97 complex. K.G.M., S.K., D.M.G., B.M., C.X. and M.R. analyzed the data. K.G.M., S.K., D.M.G. and M.R. wrote the manuscript.

## Acknowledgements

We thank all members of our laboratory for offering their ideas, help, and enthusiasm. We are grateful to the Cancer Research Laboratory Flow Cytometry Facility and the Vincent J. Proteomics/Mass Spectrometry Laboratory (NIH S10RR025622). KGM and DA were supported by NIH F32 postdoctoral fellowships; SK received support by an NSF predoctoral fellowship; and DLH is supported by a Helen Hay Whitney postdoctoral fellowship. MR is an investigator of the HHMI.

## STAR Methods

### Key resources table

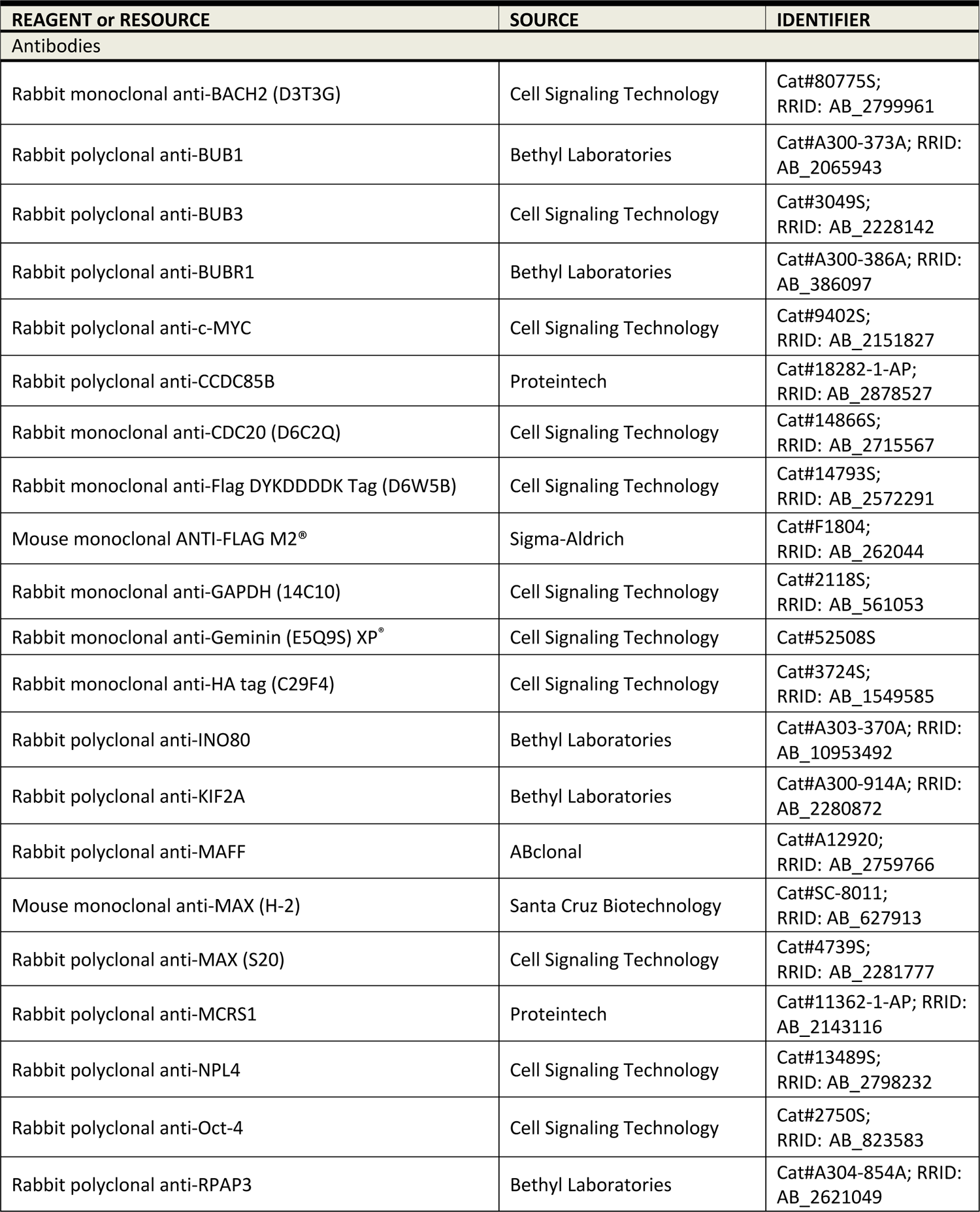
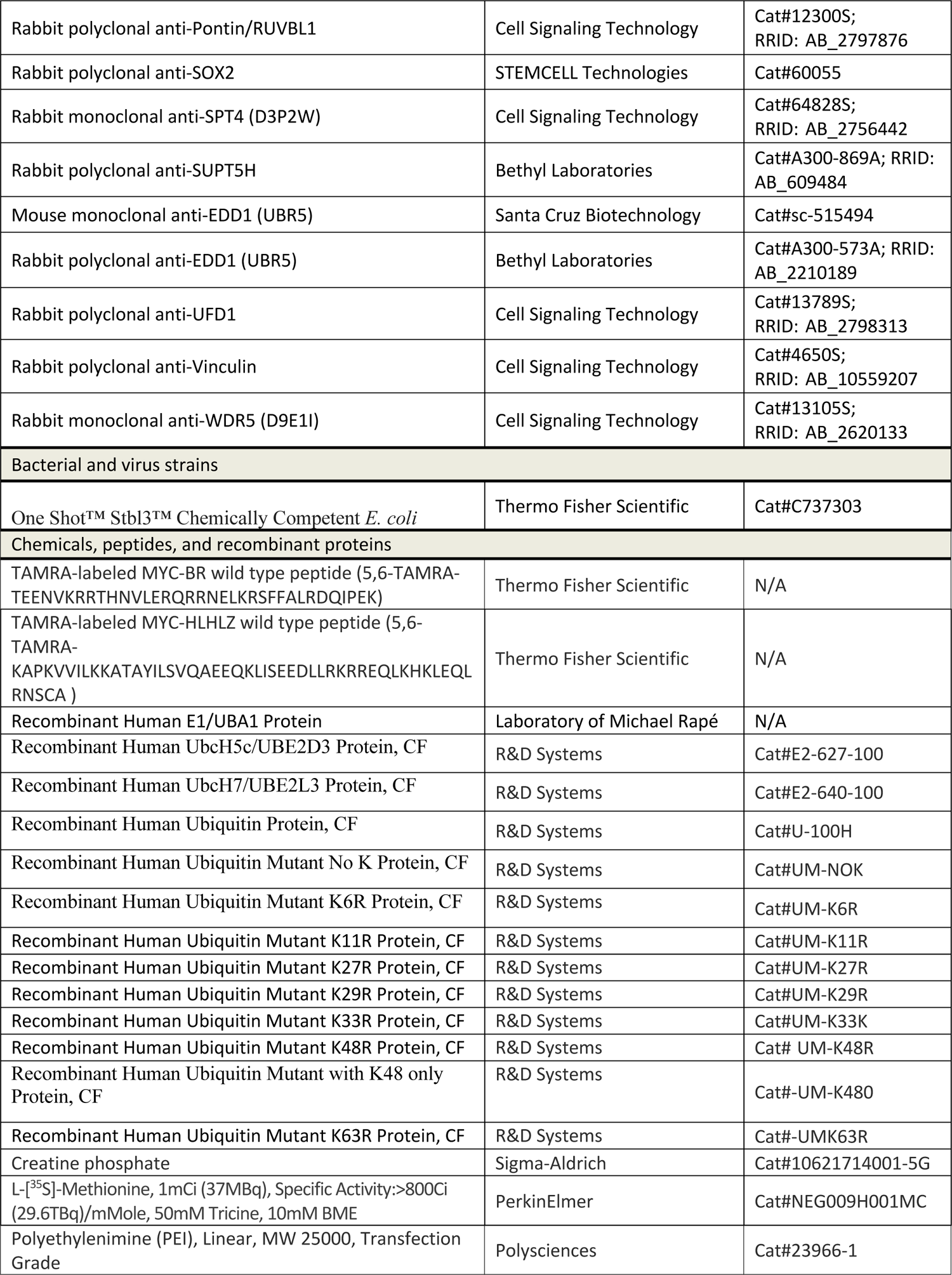
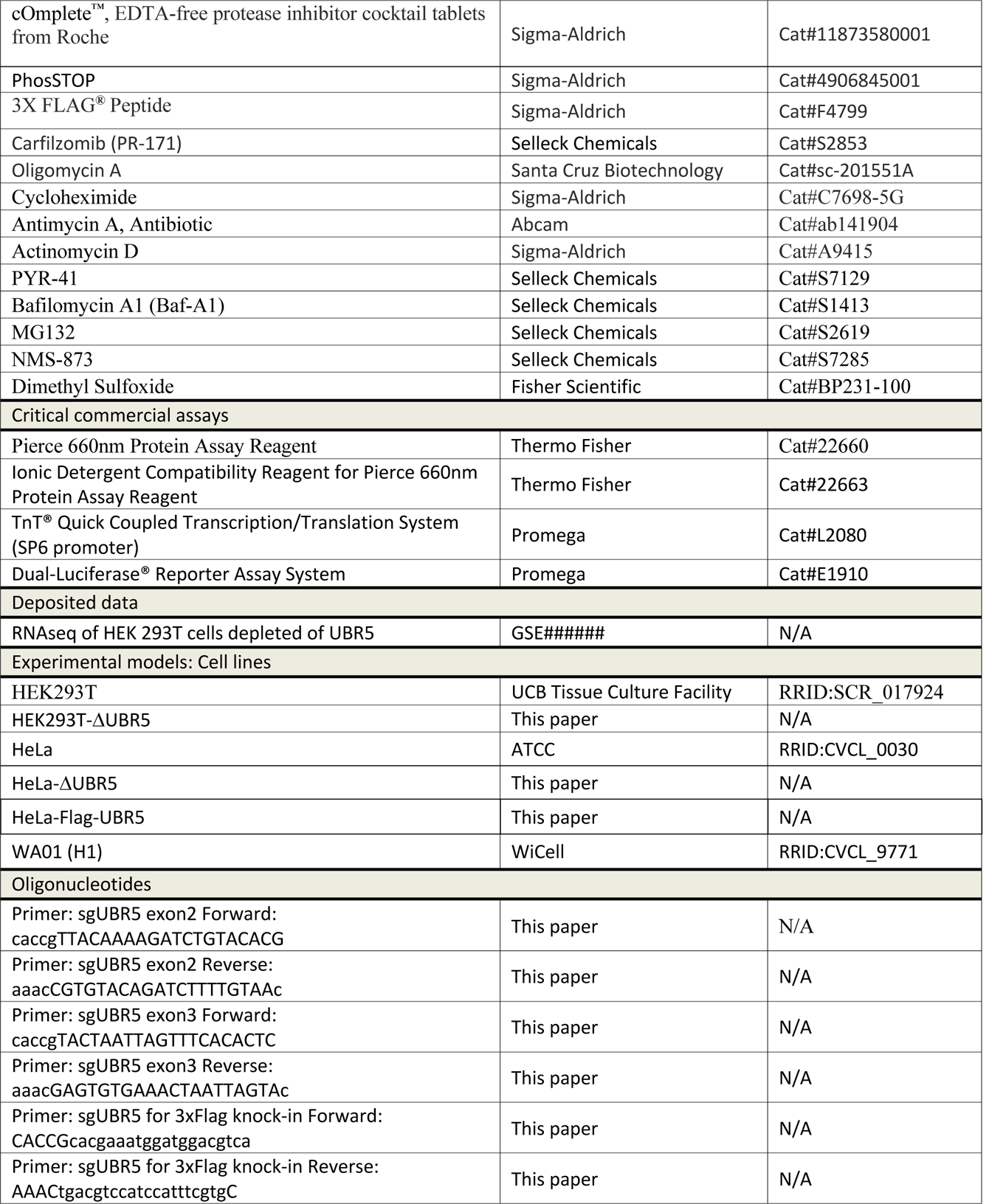
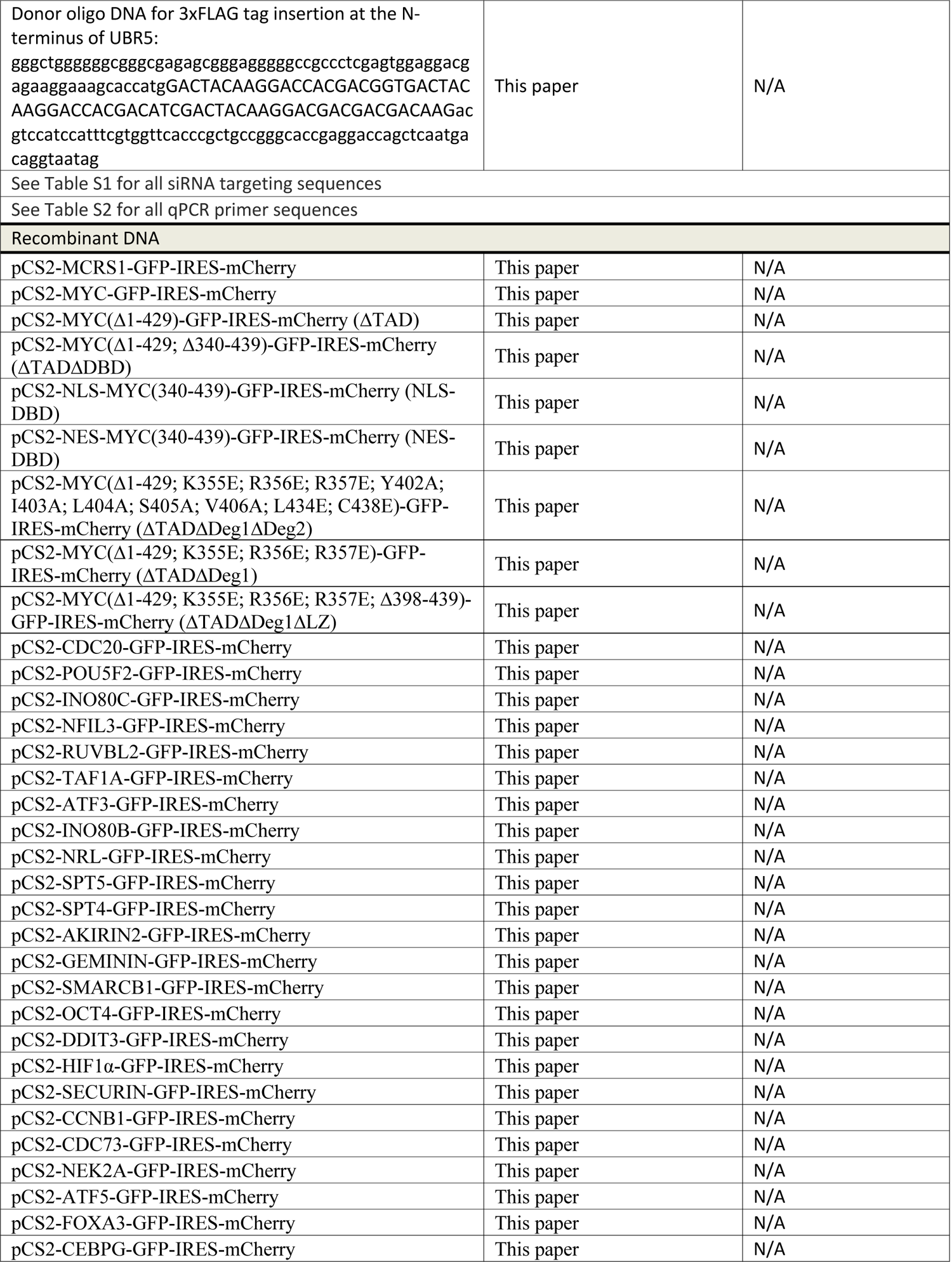
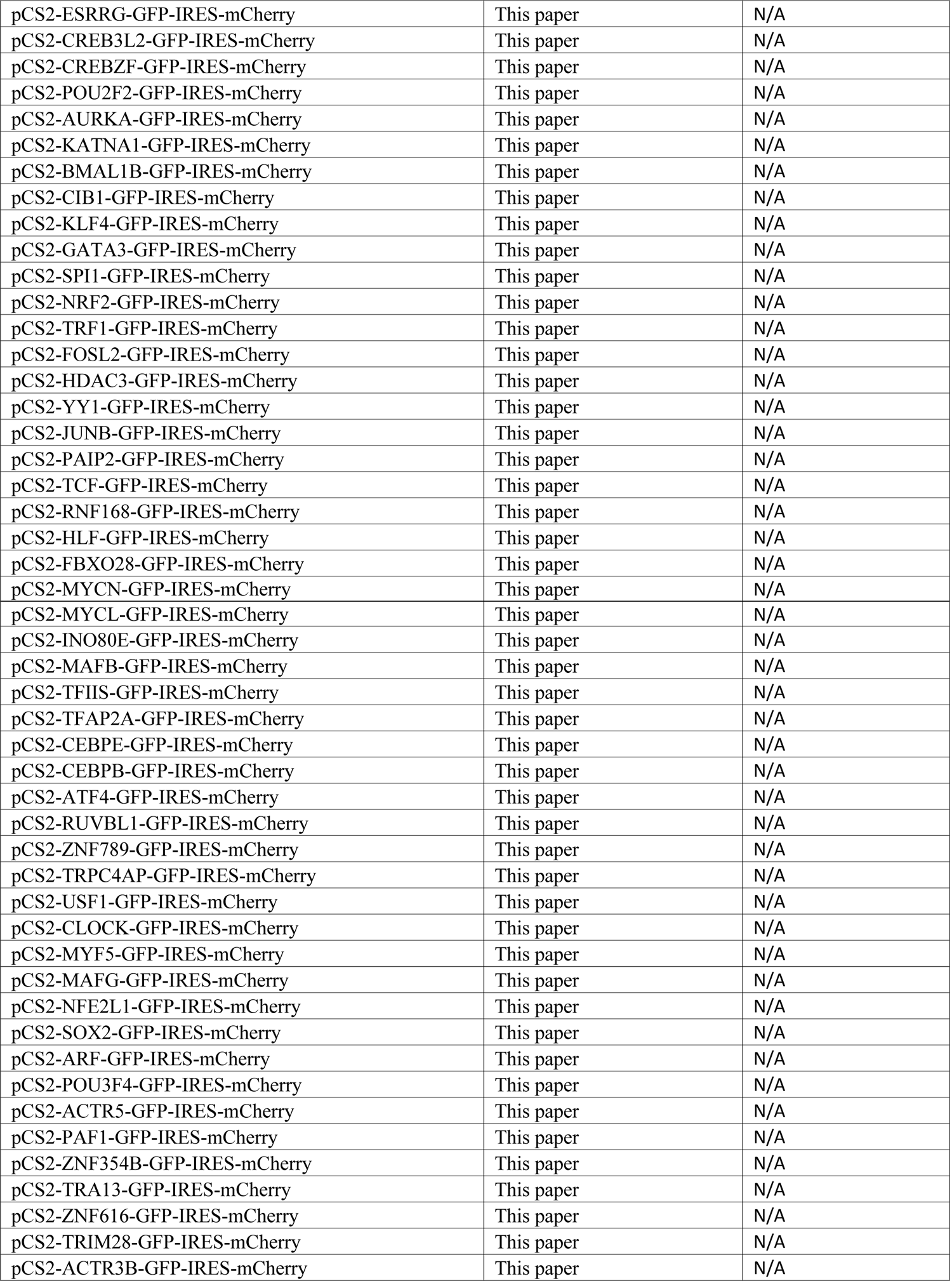
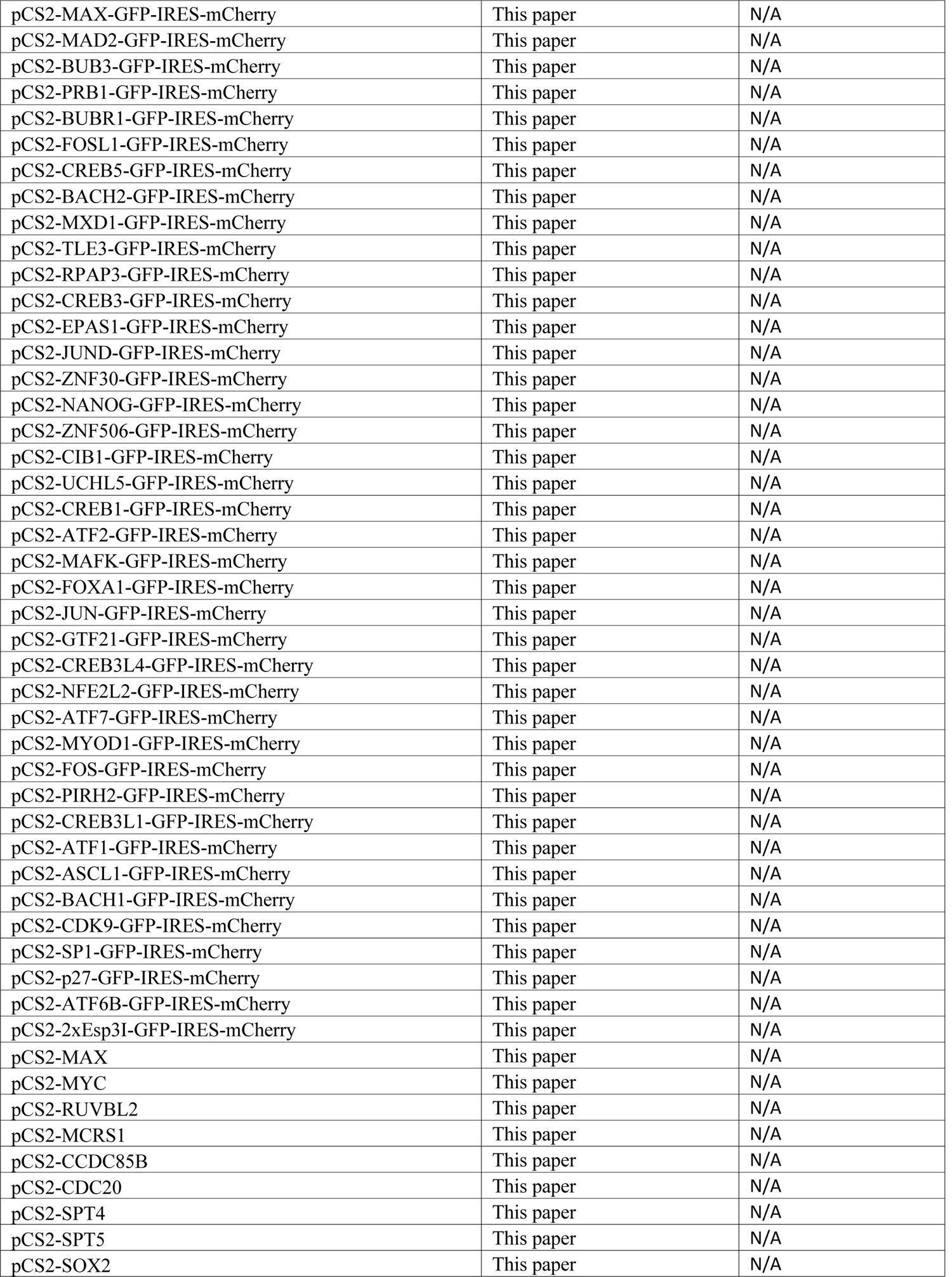
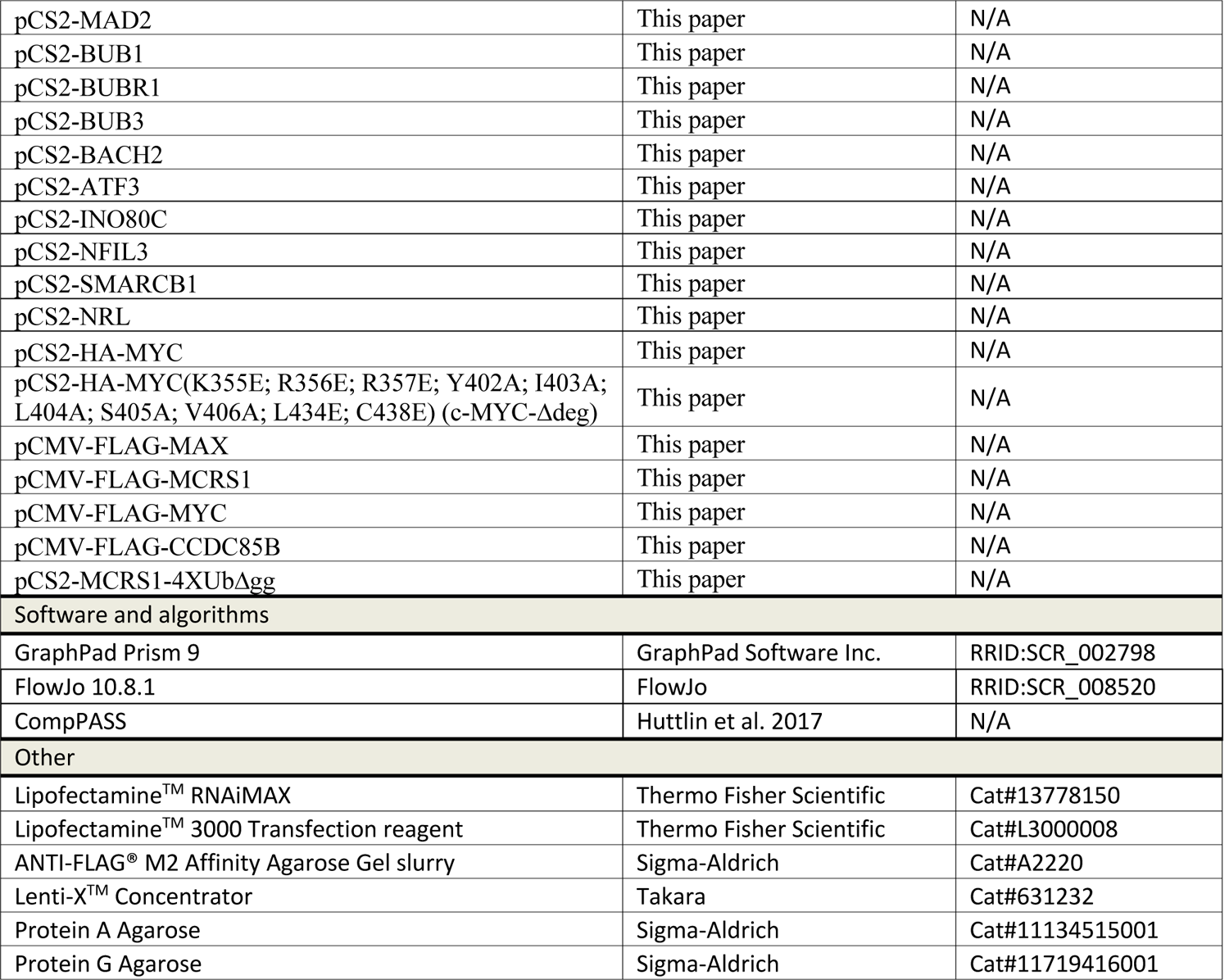

### Resource availability

#### Lead Contact

Further information and requests for reagents and resources should be directed to Michael Rapé.

#### Materials Availability

All plasmids and cell lines generated in this work can be requeted from the lead contact’s lab. All antibodies, chemicals, and most cell lines used in this study are commercially available.

#### Data and Code Availability

Original gene expression data obtained by RNA-seq from HEK 293T cells lacking UBR5 (and corresponding controls) were uploaded to GSE######.

### Mammalian cell culture

Human embryonic kidney (HEK) 293T and HeLa cells were maintained in DMEM + GlutaMAX (Gibco, 10566-016) supplemented with 10% fetal bovine serum (VWR, 89510-186). Plasmid transfections for immunoprecipitations were performed using polyethylenimine (PEI) at a 1:3 ratio of DNA (in μg) to PEI (in μl at a 1 mg ml^-1^ stock concentration). siRNA transfections were performed using 20 nM of indicated siRNAs and 5ul of RNAiMAX transfection reagent (Thermo Fisher, 13778150) per well in a 6-well cell culture plate. Lentiviruses were produced in HEK 293T cells by co-transfection of lentiviral and packaging plasmids using Lipofectamine® 3000 transfection reagent (Thermo, L3000015). Viruses were harvested 48 h post transfection, concentrated using the Lenti-X concentrator (Takara, 631232), aliquoted, and stored at −80°C for later use. HEK 293T cells were purchased directly from the Berkeley Cell Culture Facility (authenticated by short tandem repeat analysis). HeLa cells were not authenticated.

Human embryonic stem cells (WiCell, WA01/H1) were grown in mTeSR™1 media (StemCell Technologies, 85850) on hESC-qualified Matrigel-coated plates (Corning, 354277) with daily media change. H1s were passaged by Accutase (StemCell Technologies, 07920) for siRNA transfections, lentiviral infections, or routine maintenance. For siRNA transfections, single cell suspensions of H1s were generated by Accutase treatment and 2–5×10^5^ cells were seeded on a Matrigel-coated well of a 6-well plate with 1.8 ml of mTeSR™1 containing 10 μM of Y-27632 (StemCell Technologies, 72308) and a 0.2 ml mixture of indicated siRNAs (at a final concentration of 20 nM) and 5ul of RNAiMAX transfection reagent buffered in Opti-MEM per well in a 6-well cell culture plate. For lentiviral infections, single cell suspensions of H1s were generated by Accutase treatment and 1.5–3×10^5^ cells were seeded on a Matrigel-coated well of a 6-well plate with 2 ml of mTeSR™1 containing 10 μM of Y-27632, polybrene (at a final concentration of 8 μg/ml), and lentiviruses produced from HEK 293T cells (see above) for 2 h. The media was immediately exchanged with 2 ml of fresh mTeSR™1 containing 10 μM of Y-27632 only. hESCs were drug-selected 24–48 h post infection.

All cell lines were routinely tested for mycoplasma contamination using the Mycoplasma PCR Detection Kit (abmGood, G238).

### Flow cytometry

HeLa cells were seeded at 300,000 cells per well in 6-well plates. siRNAs were reverse transfected (at the time of seeding cells) with 20 nM of indicated siRNAs and 5ul of RNAiMAX transfection reagent (Thermo Fisher, 13778150) per well in a 6-well cell culture plate. The next day, 0.33 μg of ^GFP^degron-IRES-mCherry reporters, 2 μg of overexpression constructs, and empty vector up to maximum of 5 μg (without co-transfection, the total was 2 μg per well) total were combined and transfected into each well using Lipofectamine 3000 (ThermoFisher, L3000008) per manufacturer’s instructions. After 24 h, cells were harvested for flow cytometry. Cells were treated with the following reagents at indicated times before harvesting: 2 μM Carfilzomib (Selleck, PR-171) for 6 h, 700 nM Bafilomycin A1 for 6 h (Selleck, S1413), 1 μM MLN-4924 (Selleck, S7109) for 6 h, or 10 μM NMS-873 (Selleck, S7285) for 6 h. At 24 h post-transfection, cells were trypsinized and centrifuged at 300×g for 5 min. Cells were resuspended in DMEM with 10% FBS and analyzed on either BD Bioscience LSR Fortessa or LSR Fortessa X20 and FlowJo.

### Cell synchronization

CRISPR/Cas9-edited ^FLAG^UBR5 HeLa cells were synchronized as previously described (Oh et al., 2020). Cells were synchronized in S phase by addition of 2 mM thymidine for 24 h, washed with 1×PBS, and harvested by scraping. To arrest cells in prometaphase, S phase-arrested cells were subsequently washed with 1XPBS to remove excess thymidine and released into fresh media (DMEM/10% FBS) for 3 h, then treated with 5 μM S-trityl-L-cysteine (Sigma, 164739) for 12–14h. Prometaphase cells were collected by vigorous pipetting and washed with 1×PBS. Cell pellets were either immediately used in immunoprecipitation assays or frozen in liquid nitrogen and stored at −80°C for later use.

### Purification of UBR5

Human UBR5 enzyme was purified from extracts of CRISPR/Cas9-edited ^FLAG^UBR5 HeLa cells. Harvested prometaphase pellets were lysed in lysis buffer (20 mM HEPES, pH 7.4, 5 mM KCl, 150 mM NaCl, 1.5 mM MgCl_2_, 0.1% Nonidet P-40, 1X cOmplete™ protease inhibitor cocktail (Roche, 04693159001), 2 μM carfilzomib (Selleck, PR-171) and 1 μl of benzonase (Millipore, 70746) per 15-cm plate). Detergent lysed cells were then subjected to a high-speed spin (20,000×*g*) to remove cellular debris and the clarified extract was pre-cleared with protein A-agarose resin (Roche, 11719408001). UBR5 was purified with anti-FLAG® M2 affinity resin (Sigma, A2220) for 1.5 h at 4°C. UBR5-coupled beads were washed 5× with lysis buffer (minus inhibitors and benzonase) prior to use.

### In vitro transcription/translation (IVT/T) of substrates

All *in vitro* synthesized substrates were cloned under the SP6 promoter. The corresponding plasmids can be found in the Star Methods table. ^35^S-labeled substrates were generated by incubating 583 ng of plasmid DNA in 14 μl of rabbit reticulocyte lysate (Promega, L2080) supplemented with 2 μM carfilzomib and 0.3 μl of ^35^S-Met (PerkinElmer, NEG009H001MC) for 1 h at 30°C. ^35^S-labeled substrates were used for *in vitro* ubiquitylation assays.

### In vitro ubiquitylation

*In vitro* ubiquitylation assays were performed in a 10 μl reaction volume: 0.5 μl of 10 μM E1 (250 nM final), 1 μl of 25 μM E2 (2.5 μM final), 1 μl of 10 mg ml^-1^ ubiquitin (1 mg ml^-1^ final) (R&D Systems, U-100H), 1 μl of 100 mM DTT, 1.5 μl of energy mix (150 mM creatine phosphate, 20 mM ATP, 20 mM MgCl_2_, 2 mM EGTA, pH to 7.5 with KOH), 1 μl of 1×PBS, 1 μl of 10× ubiquitylation assay buffer (250 mM Tris 7.5, 500 mM NaCl, and 100 mM MgCl_2_), and 3 μl of substrate (*in vitro* translated or recombinant) were pre-mixed and added to 10 μl of UBR5-coupled bed resin (see section on Purification of UBR5). Reactions were performed at 30°C with shaking for 2 h unless noted otherwise. Reactions were stopped by adding 2X urea sample buffer and resolved on SDS-acrylamide gels prior to autoradiography. E1 was purified as described in Meyer and Rape (2014) while commercially available UBE2D3 (R&D Systems, E2-627-100) and UBE2L3 (R&D Systems, E2-640-100) were used.

### Mass spectrometry

Mass spectrometry was performed on immunoprecipitates prepared from HEK 293T or HeLa cells. For immunoprecipitations of overexpressed proteins, thirty 15-cm plates of HEK 293T cells were PEI-transfected, grown to confluence, harvested, and lysed in lysis buffer (20 mM HEPES, pH 7.4, 5 mM KCl, 150 mM NaCl, 1.5 mM MgCl_2_, 0.1% Nonidet P-40, and 1× cOmplete™ protease inhibitor cocktail). For endogenous UBR5 IPs, one hundred 15-cm plates of CRISPR/Cas9-edited ^FLAG^UBR5 HEK 293T or HeLa cells were used. Lysed extracts were clarified by high-speed centrifugation, pre-cleared with protein A-agarose slurry and bound to anti-FLAG® M2 affinity resin. IPs were then washed and eluted 3× at 30°C with 0.5 mg ml^-1^ of 3×FLAG® peptide (Sigma, F4799) buffered in 1×PBS plus 0.1% Triton X-100. Elutions were pooled and precipitated overnight at 4°C with 20% trichloroacetic acid. IPs were then pelleted, washed 3× with an ice-cold acetone/0.1 N HCl solution, dried, resolubilized in 8 M urea buffered in 100 mM Tris 8.5, reduced with TCEP (at a final concentration of 5 mM) for 20 min, alkylated with iodoacetamide (at a final concentration of 10 mM) for 15 min, diluted four-fold with 100 mM Tris 8.5, and digested with 0.5 mg ml^-1^ of trypsin supplemented with CaCl_2_ (at a final concentration of 1 mM) overnight at 37°C. Trypsin-digested samples were submitted to the Vincent J. Coates Proteomics/Mass Spectrometry Laboratory at UC Berkeley for analysis. Peptides were processed using multidimensional protein identification technology (MudPIT) and identified using a LTQ XL linear ion trap mass spectrometer. To identify high confidence interactors, CompPASS analysis of the query mass spectrometry result was performed against mass spectrometry results from unrelated FLAG immunoprecipitates performed in our laboratory.

### FLAG immunoprecipitation

Overexpression constructs were PEI-transfected into HEK 293T or wild-type HeLa cells with 10 μg of each plasmid per 15-cm plate for 48 h prior to harvesting. Human ^FLAG^UBR5 complexes were immunoprecipitated from endogenously FLAG-tagged HeLa cell lines grown on 15-cm plates to near-confluency. Cells were harvested by scraping and pellets were lysed in lysis buffer (20 mM HEPES, pH 7.4, 5 mM KCl, 150 mM NaCl, 1.5 mM MgCl_2_, 0.1% Nonidet P-40, 1× cOmplete™ protease inhibitor cocktail (Roche, 04693159001), and 1 μl of benzonase (Millipore, 70746) per 15-cm plate). Detergent lysed cells were then subjected to a high speed spin (20,000 × *g*) to remove cellular debris and the clarified extract was pre-cleared with protein A-agarose resin (Sigma-Aldrich, 11134515001). Bait proteins were purified with anti-FLAG® M2 affinity resin (Sigma, A2220) for 1.5 h at 4°C. For *in vitro* ubiquitylation reactions, ^FLAG^UBR5-coupled beads were washed 5× with lysis buffer (minus inhibitors and benzonase) prior to use. For western blots, FLAG beads with captured bait protein were eluted at 30°C with 0.5 mg ml^-1^ of 3×FLAG® peptide (Sigma, F4799) buffered in 1×PBS plus 0.1% Triton X-100 for 20 min, shaking. Eluates were combined with 2× urea sample buffer (120 mM Tris pH 6.8, 4% SDS, 4 M urea, 20% glycerol, bromophenol blue) prior to SDS-PAGE.

### Purification of p97, Ufd and Npl4

Human p97 was subcloned into pET28a His-tagged expression vector (pET28a-6xHis-FLAG-TEV-p97) and were expressed in LOBSTR-BL21(DE3)-RIL competent cells. Human Ufd1 and Npl4 were subcloned into a pET28a His-tagged expression vector (pET28a-Ufd1-6xHis) and a pMAL expression vector (pMAL-Npl4) respectively, and co-expressed in LOBSTR-BL21(DE3)-RIL competent cells. Protein expression was induced at log phase with 500 µM IPTG for 16 hours at 18°C. Cells were lysed in Lysis buffer (100mM Tris pH 7.4, 500 mM KCl, 5 mM MgCl_2_, 20 mM imidazole, 5 mM BME, 5% glycerol, and protease inhibitors) using a LM10 Microfluidizer. Lysate was clarified prior to a 1 hour incubation with equilibrated Ni-NTA agarose beads (Qiagen, Cat# 20350), and beads were washed in wash buffer (50 mM HEPES pH 7.4, 150mM KCl, 5 mM MgCl_2_, 5 mM BME, 20mM imidazole and 2.5% glycerol). p97 was eluted in wash buffer containing 250 mM imidazole.

Imidazole-eluted fractions containing p97, Ufd1 or Npl4 were confirmed by Coomassie-stained SDS-PAGE were subject to further separation on a HiLoad 26/600 Superdex 200 size exclusion chromatography column (Cytiva, Cat# 28989336) in SEC buffer (20 mM HEPES, pH 7.4, 150 mM KCl, 1mM MgCl_2_, and 2.5% glycerol) and peak fractions were collected and concentrated with Centricon® Plus-70 Centrifugal Filter Units (Millipore, Cat# UFC703008), filter sterilized, and snap frozen in PBS at −80°C.

### Pulldown of ubiquitylated substrate by p97-Ufd1-Npl4 complexes

*In vitro* ubiquitylation reactions were performed as described in the relevant section. Three reactions of ubiquitylated substrate were used per pulldown. Purified p97, Ufd1, and Npl4 were diluted to 8 μg ml^-1^ in binding buffer (20 mM HEPES pH 7.4, 150 mM KCl, 1mM MgCl_2_, 0.05% NP-40) and incubated with ubiquitylated material for 1 h at 4°C. Complexes were combined with 10 μl of anti-FLAG® M2 affinity resin (Sigma, A2220) for 1.5 h at 4°C. Beads were washed five times with binding buffer and resuspended in 2× urea sample buffer (120 mM Tris pH 6.8, 4% SDS, 4 M urea, 20% glycerol, bromophenol blue) prior to SDS-PAGE and autoradiography.

### Real-time qPCR (qRT-PCR) analysis

For qRT–PCR analysis, total RNA was purified from cells using the NucleoSpin® RNA kit (Macherey-Nagel, no. 740955). For each sample, 1 μg of total RNA was reverse transcribed using the RevertAid First Strand cDNA Synthesis kit (ThermoFisher, K1622) and then diluted 10-fold for qRT-PCR. Expression levels were quantified using the Roche KAPA SYBR FAST qPCR Kit (Roche, KK4602) on a Roche LightCycler® 480 II. qRT-PCR primers used in this study can be found in Table S2.

### Generation of CRISPR/Cas9 genome-edited cell lines

All cell lines used in this publication were generated from HEK 293T cells or HeLa cells. The <u>guide</u> <u>RNA</u> sequences were designed using the online resource provided by the Zhang Lab at MIT (http://crispr.mit.edu). The sequences of the genes for editing were obtained from the UCSC Genome browser. The oligonucleotides for guide RNAs (listed in the <u>Key Resources Table</u>) and their complementary sequences were ordered from IDT, annealed, and cloned into a pX330 vector according to the protocol at https://benchling.com/protocols/5DmqRd/crispr-mediated-gene-disruption-in-ch12f3-2-cells/sbs. For CRISPR/Cas9-mediated <u>gene disruption</u>, HEK 293T cells cultured in a 6-well plate at 50% confluence were transfected using TransIT-293 transfection reagent (Mirus) with two pX330 plasmids (2 μg of total DNA) each encoding a guide RNA for site-specific gene cutting. The guides for cutting were designed to create deletions and introduce open reading frame (ORF) shifts for gene disruption. Three days post-transfection, a sample of transfected cells was treated with DNA extraction solution (QuickExtract, Epicenter) and editing was assessed by PCR amplification of the sequence of interest using specific PCR primers. Clonal selection was performed by seeding the cells into 96-well plates at one cell per well density, allowing the single cells to expand, and checking editing by PCR. The clones were validated using western blotting with specific antibodies.

### Guide RNAs for CRISPR/Cas9-mediated genome editing

A pair of RNA guides was used to remove part of the second and third exons of the *UBR5* locus in HEK 293T cells and HeLa cells: TTACAAAAGATCTGTACACG (Exon 2), TACTAATTAGTTTCACACTC (Exon 3).

### RNAseq sample preparation and analysis

Wild-type and UBR5 knockout HEK 293T cells were grown in a 6-well plate and collected for transcriptomic analysis. For each sample, there are three biological replicates. Total RNA was extracted from cells using the NucleoSpin® RNA kit (Macherey-Nagel, no. 740955) according to the manufacturer’s instructions. Library preparation and sequencing was executed by Novogene Bioinformatics Technology Co., Ltd. (Beijing, China).

Genes identified by RNAseq were subjected to bioinformatic analysis. Transcription factor binding sites were identified using the UCSC_TFBS feature of DAVID 2021, a web-based application (https://david.ncicrf.gov).

## Supplemental Figures

**Figure S1:**
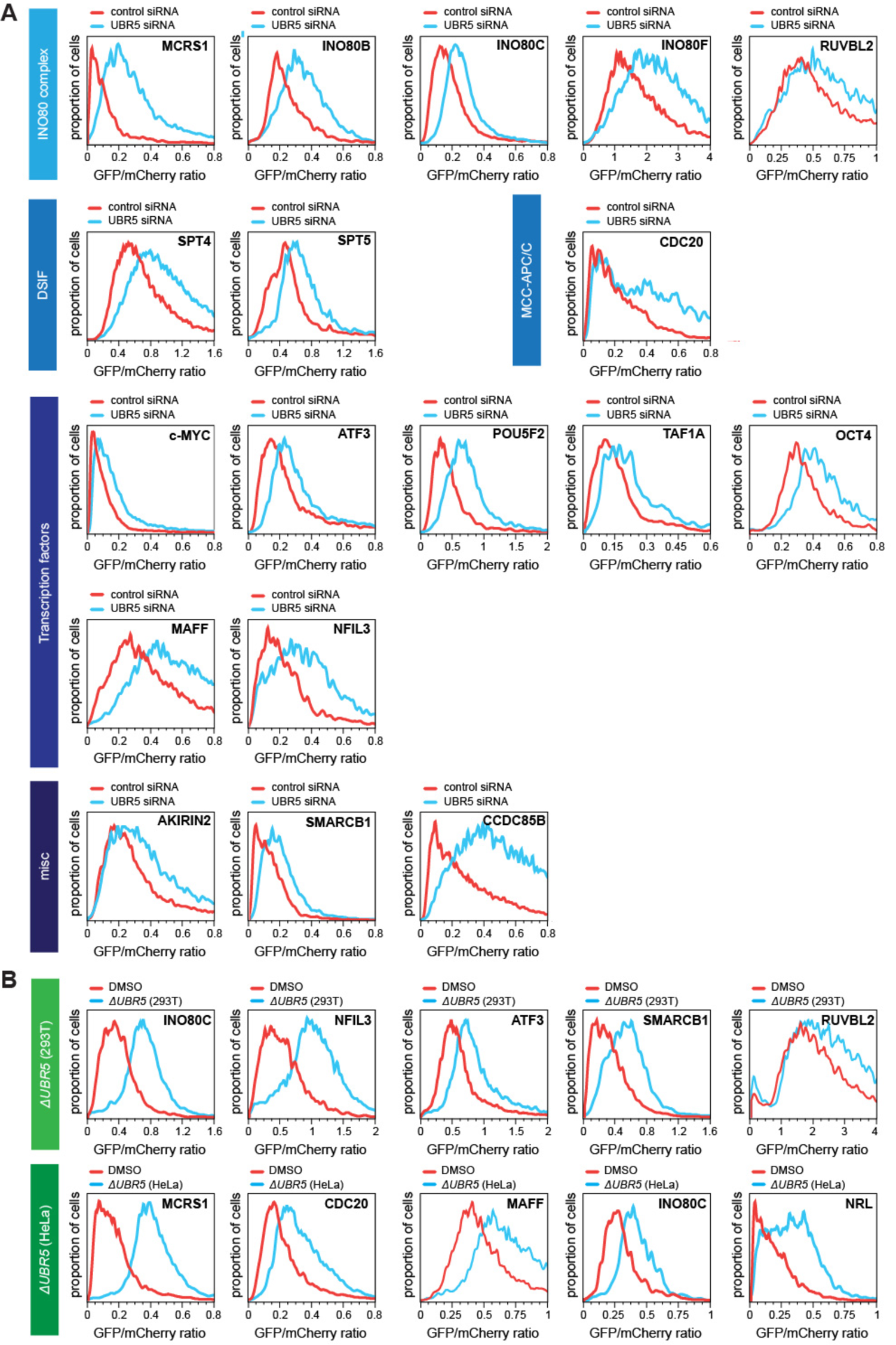
Validation of UBR5 substrate screen. **A.** Validation of UBR5 substrate screen in cells treated with either control siRNA or siRNAs targeting UBR5. Substrate stability was measured as a GFP-tagged protein co-expressed with mCherry by FACS. **B.** Validation of UBR5 screen for select substrates in *ΔUBR5* (293T) and *ΔUBR5* (HeLa) cells. Substrate stability was measured by FACS, as described above.

**Figure S2:**
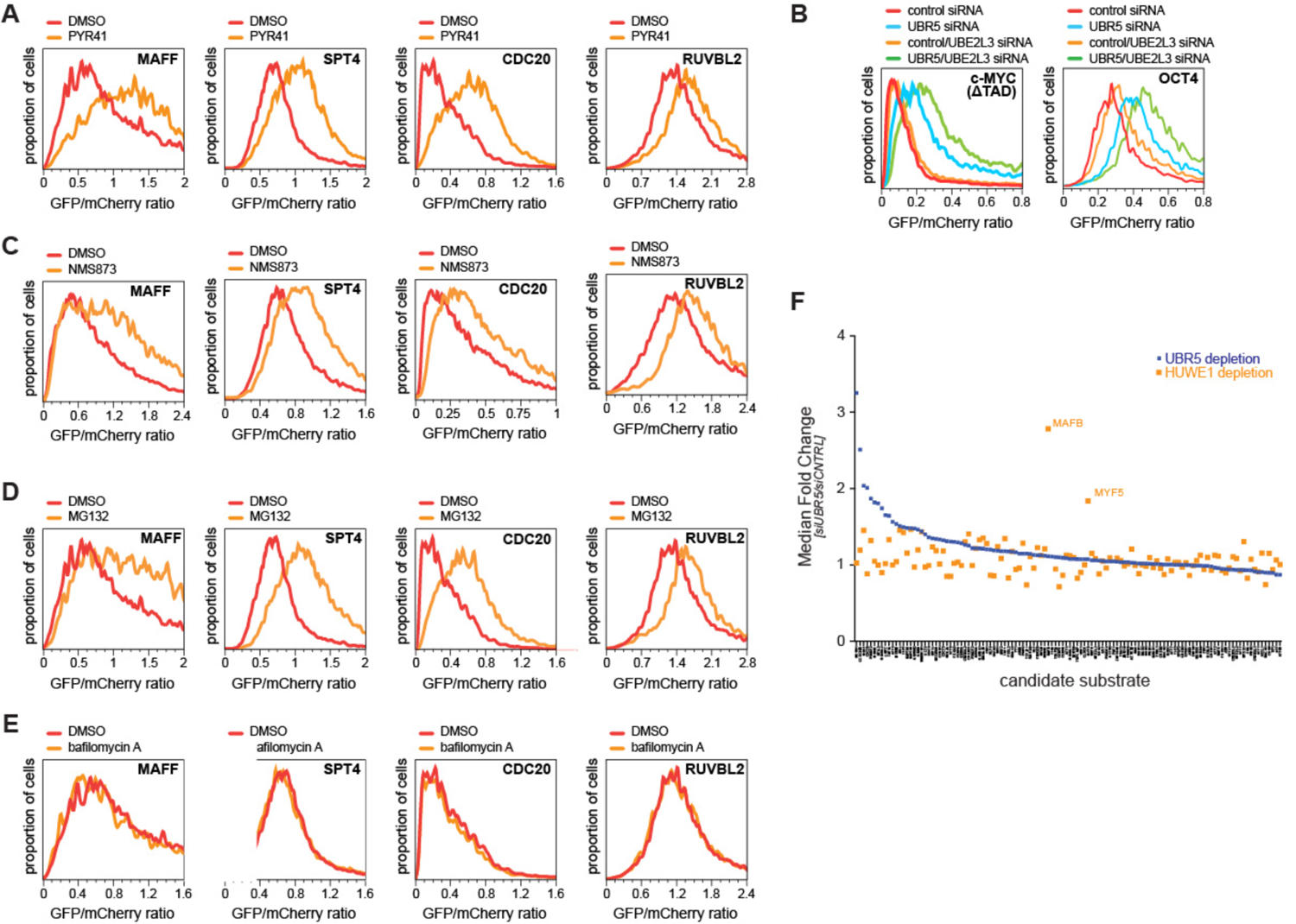
UBR5 substrates are subject to ubiquitin-, proteasome-, and p97-dependent degradation. **A.** Select substrates are stabilized upon inhibition of the E1 activating enzyme by PYR41. Substrate stability was measured using the GFP/mCherry reporter by FACS. **B.** UBR5 cooperates with UBE2L3 to degrade substrates. Cells were treated with control siRNAs, either UBR5 or UBE2L3 siRNAs, or with both UBR5 and UBE2L3 siRNAs. Substrate stability was measured using the GFP/mCherry reporter by FACS. **C.** Select substrates are stabilized by inhibition of p97/VCP with NMS873. Substrate stability was measured using the GFP/mCherry reporter by FACS. **D.** Select substrates are stabilized by proteasome inhibition with MG132. Stability was measured using the GFP/mCherry reporter by FACS. **E.** Lysosome inhibition by bafilomycin A does not affect substrate stability, as measured using the GFP/mCherry reporter by FACS. **F.** Focused stability screen to test for effects of HUWE1 depletion. Candidate substrates of our UBR5 target screen were expressed in cells treated with control siRNA or siRNAs targeting UBR5 as GFP-tagged reporters together with mCherry. Protein abundance was measured as the ratio between GFP and mCherry, as determined by FACS.

**Figure S3:**
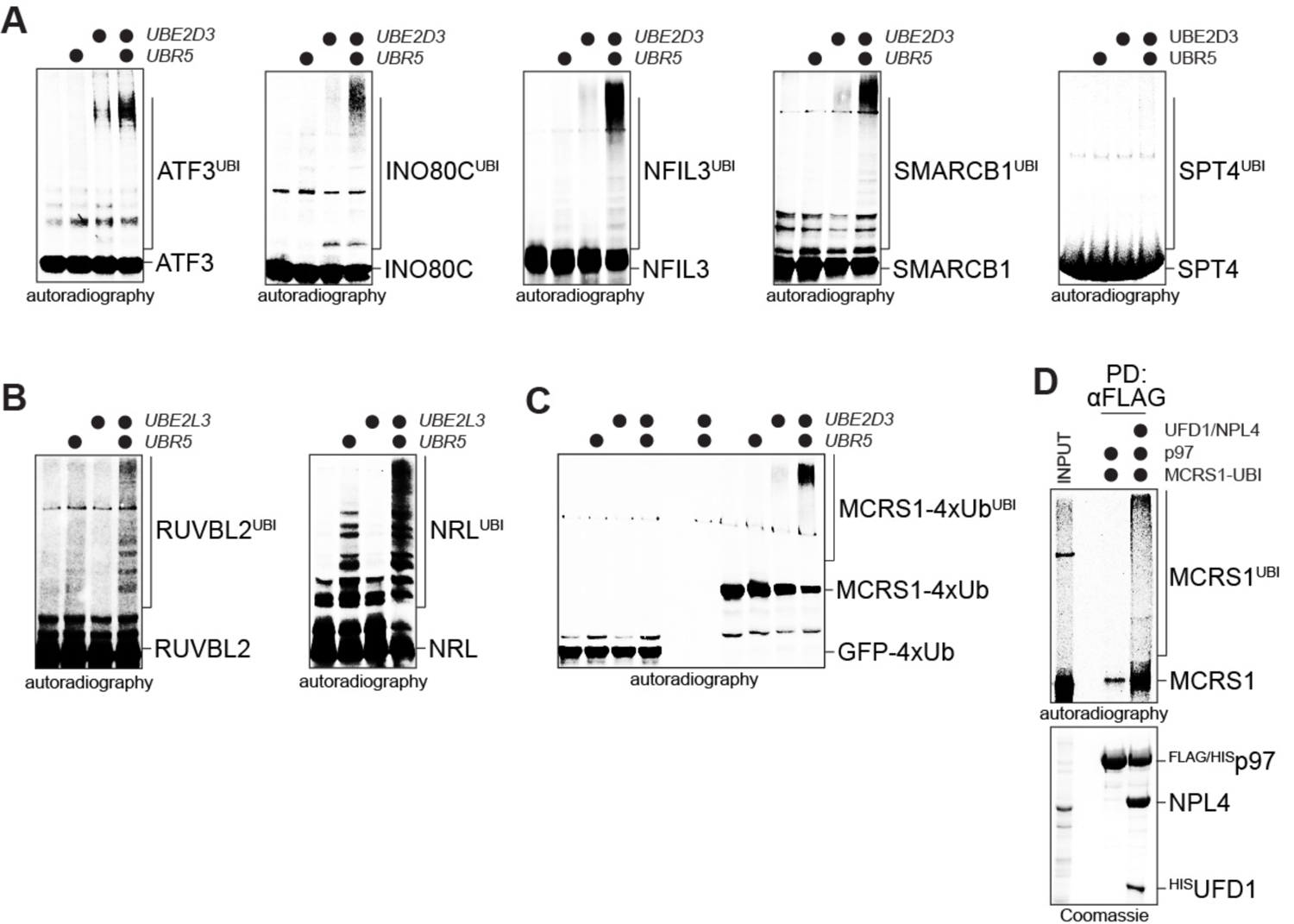
UBR5 ubiquitylates transcriptional regulators. **A.** *In vitro* ubiquitylation of candidate substrates by UBR5 and UBE2D3. Targets were produced as ^35^S-labeled proteins by IVT/T and incubated with UBR5 and recombinant E1, UBE2D3, and ubiquitin. Target modification was visualized by autoradiography. **B.** UBR5 can collaborate with UBE2L3 *in vitro*. ^35^S-labeled targets were incubated with UBR5, E1, UBE2L3, and ubiquitin and analyzed as above. **C.** Chain initiation greatly enhances UBR5 activity. GFP∼Ub4 (four ubiquitin moieties fused to GFP) or MCRS1∼Ub4 were incubated with UBR5, E1, UBE2D3, and ubiquitin and analyzed for ubiquitylation as described above. **D.** Ubiquitylated MCRS1 is recognized by p97. Purified complexes of recombinant p97 and its adaptors UFD1 and NPL4 were immobilized and incubated with MCRS1 that had been ubiquitylated by UBR5 and UBE2D3 *in vitro*. Retention of ubiquitylated MCRS1 is detected by autoradiography.

**Figure S4:**
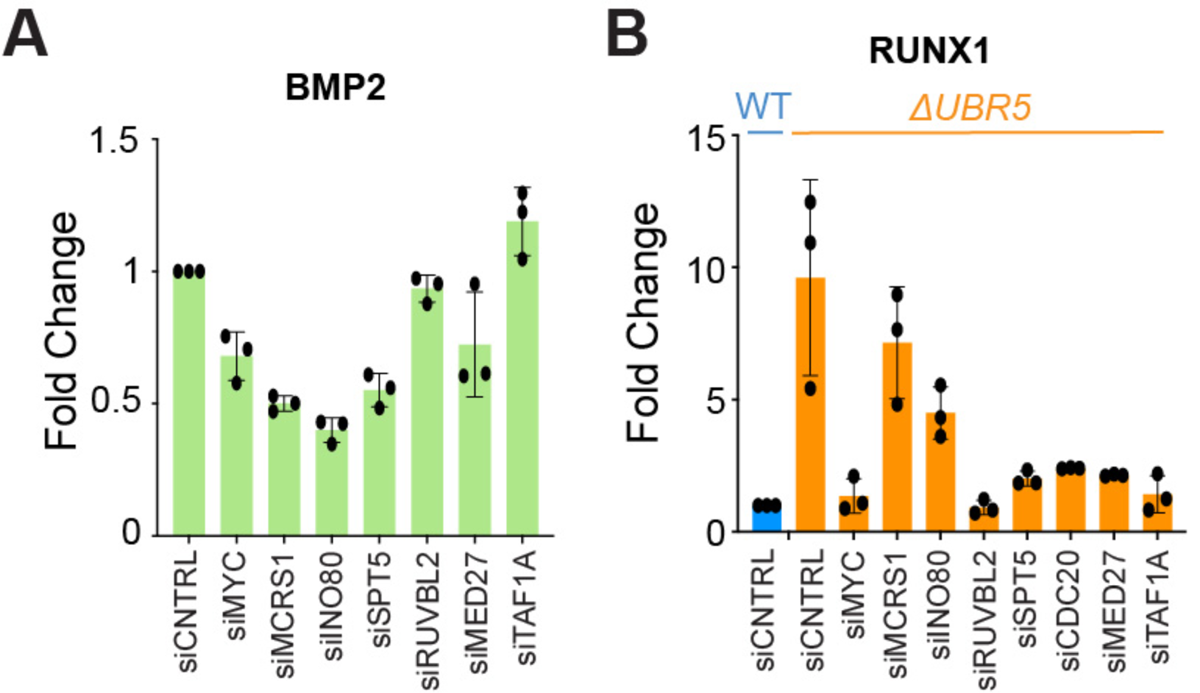
UBR5 targets are required for accurate gene expression. **A.** Partial depletion of UBR5 substrates and network components reduces expression of the UBR5-dependent gene *BMP2*, as determined by qRT-PCR. **B.** Partial depletion of UBR5 substrates and network components blunts the effects of *UBR5* deletion onto expression of RUNX1, as determined by qRT-PCR.

**Figure S5:**
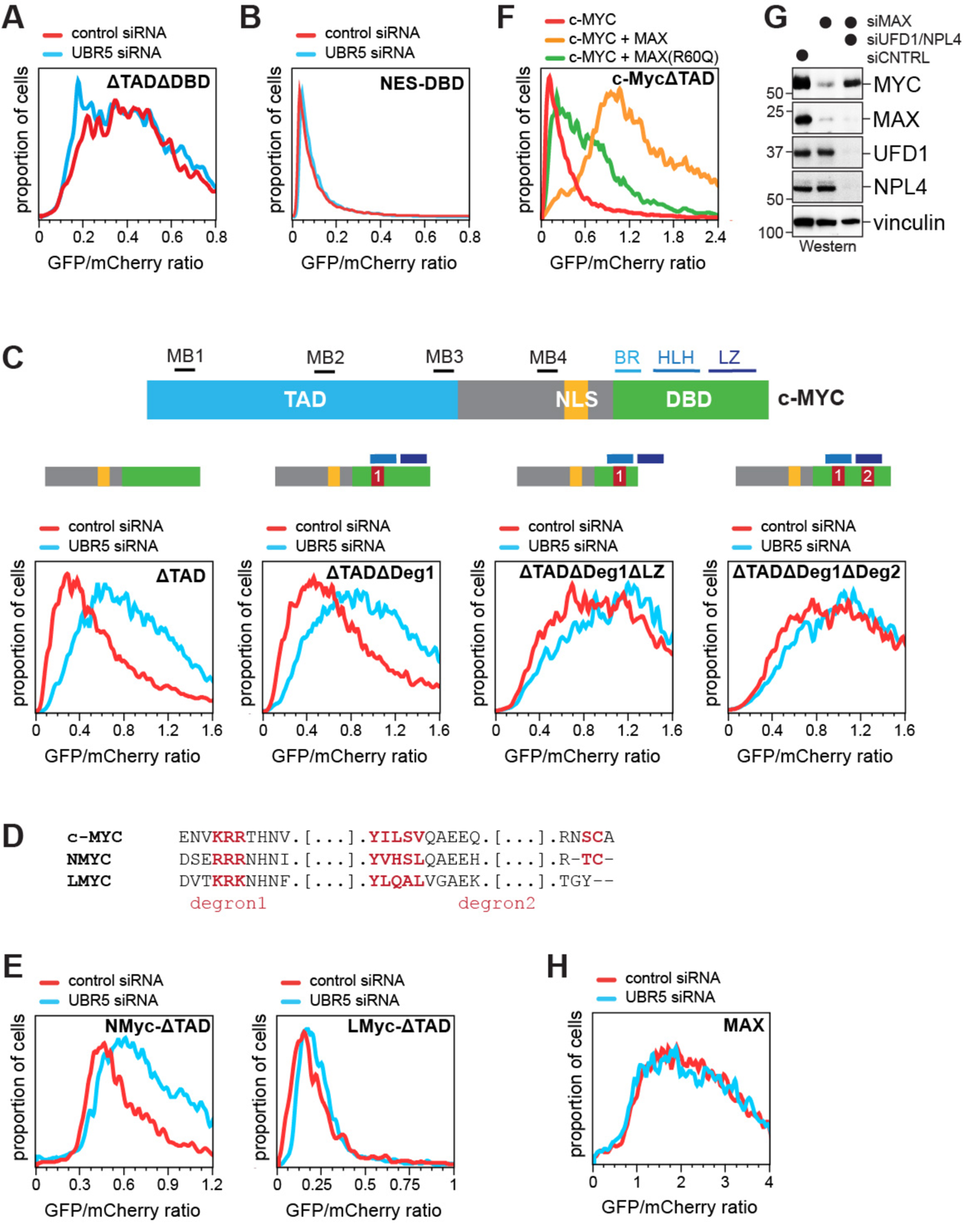
UBR5 targets orphan c-MYC for degradation. **A.** The DNA-binding domain of c-MYC (DBD) is required for UBR5-dependent degradation. A GFP-c-MYC^ΔTADΔDBD^::mCherry reporter was expressed in cells treated with control or UBR5 siRNA, and protein stability was measured by FACS. **B.** The DNA-binding domain of c-MYC (DBD) is degraded in the cytoplasm independently of UBR5. A GFP-c-MYC^DBD^ reporter fused to a strong nuclear export signal (NES) was expressed in cells treated with control or UBR5 siRNAs and protein stability was measured by FACS. **C.** c-MYC contains two redundant degrons for UBR5 in its DNA-binding domain. GFP-tagged reporters containing the carboxy-terminal half of c-MYC (ΔTAD); mutations in degron 1 (ΔTADΔDeg1); additional deletion of the leucine zipper (ΔTADΔDeg1ΔLZ); or mutation of both degrons (ΔTADΔDeg1ΔDeg2) were expressed in cells treated with control siRNA or siRNAs targeting UBR5. Protein stability was measured by FACS. **D.** Conservation of degron 1 and degron 2 (consisting of both red stretches) between c-MYC, NMYC, and LMYC. **E.** NMYC, but not LMYC, is a UBR5 substrate. Reporters containing NMYC-ΔTAD or LMYC-ΔTAD were expressed in cells treated with control siRNA or siRNAs targeting UBR5. Protein stability was measured by FACS. **F.** A MAX^R60Q^ mutant, which is less efficient in forming dimers with c-MYC, is also less efficient in stabilizing a c-MYC^ΔTAD^ reporter than wildtype MAX, as determined by FACS. **G.** Co-depletion of the p97 adaptors UFD1 and NPL4 rescues c-MYC levels in cells depleted of MAX, as determined by Western blotting. **H.** MAX is not degraded by UBR5, as determined by monitoring stability of a GFP-MAX reporter using FACS.

**Figure S6:**
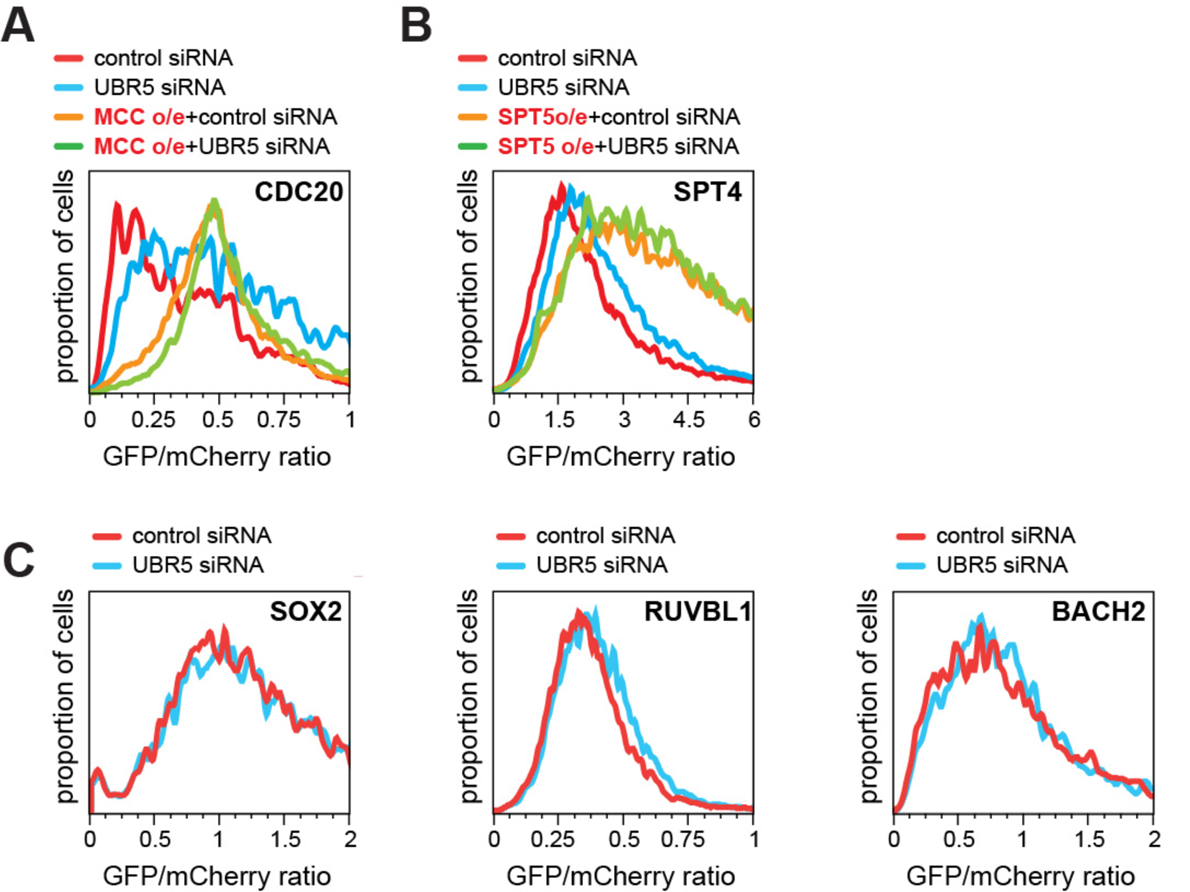
Complex formation protects URB5 targets from degradation. **A.** Binding of CDC20 by MCC components (BUBR1, BUB3, MAD2) protects a GFP-CDC20::mCherry reporter from degradation, as determined by FACS. **B.** A SPT4 reporter is stabilized more extensively upon co-expression of SPT5 than by depletion of UBR5, as determined by FACS. **C.** The stabilizing partners SOX2, RUVBL1, and BACH2, are not significantly stabilized by UBR5 depletion, as determined by FACS.

**Figure S7:**
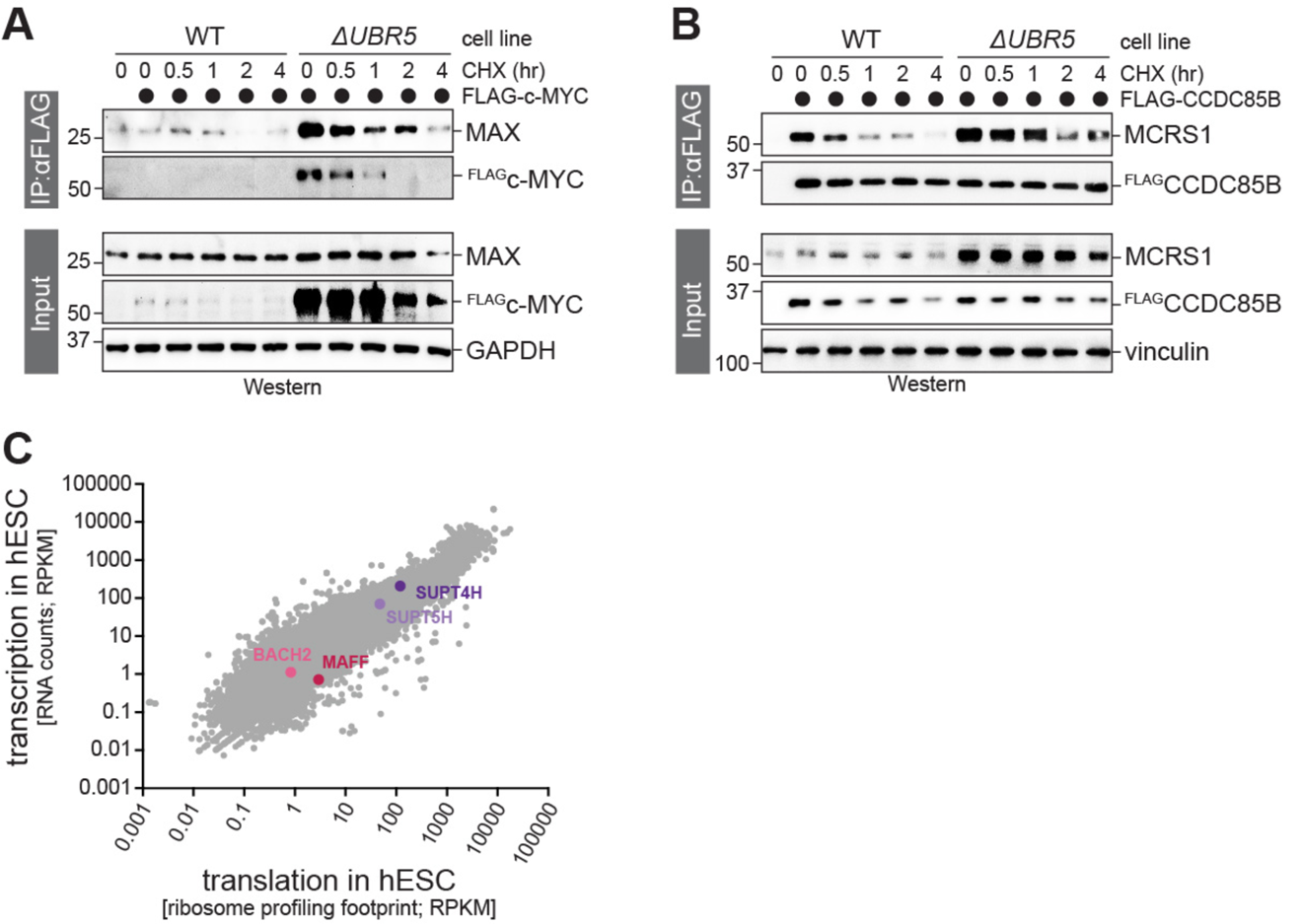
Orphan protein degradation regulates complex stability. **A.** *UBR5* deletion leads to a strong accumulation of transiently expressed c-MYC and c-MYC/MAX complexes. Control or *ΔUBR5* cells were transfected with ^FLAG^c-MYC and treated with cycloheximide as indicated. ^FLAG^c-MYC was immunoprecipitated and co-purifying MAX was detected by Western blotting. **B.** *UBR5* increases the persistence of CCDC85B/MCRS1 complexes after inhibition of mRNA translation. Control or *ΔUBR5* cells were transfected with ^FLAG^CCDC85B and treated with cycloheximide as indicated. ^FLAG^CCDC85B was immunoprecipitated and co-purifying MCRS1 was detected by Western blotting. **C.** hESCs produce unstable complex subunits in excess over their stable binding partners. Combined RNAseq and ribosome profiling experiments in hESCs show increased expression of MAFF over BACH2 and SPT5 over SPT4.

## References

1. Adhikary, S., Marinoni, F., Hock, A., Hulleman, E., Popov, N., Beier, R., Bernard, S., Quarto, M., Capra, M., Goettig, S., et al. (2005). The ubiquitin ligase HectH9 regulates transcriptional activation by Myc and is essential for tumor cell proliferation. Cell 123, 409–421.

2. Ali, A., Veeranki, S.N., Chinchole, A., and Tyagi, S. (2017). MLL/WDR5 Complex Regulates Kif2A Localization to Ensure Chromosome Congression and Proper Spindle Assembly during Mitosis. Dev Cell 41, 605–622 e607.

3. Ali, I., Ruiz, D.G., Ni, Z., Johnson, J.R., Zhang, H., Li, P.C., Khalid, M.M., Conrad, R.J., Guo, X., Min, J., et al. (2019). Crosstalk between RNA Pol II C-Terminal Domain Acetylation and Phosphorylation via RPRD Proteins. Mol Cell 74, 1164–1174 e1164.

4. Aoi, Y., Takahashi, Y.H., Shah, A.P., Iwanaszko, M., Rendleman, E.J., Khan, N.H., Cho, B.K., Goo, Y.A., Ganesan, S., Kelleher, N.L., et al. (2021). SPT5 stabilization of promoter-proximal RNA polymerase II. Mol Cell 81, 4413–4424 e4415.

5. Apostolou, E., and Stadtfeld, M. (2018). Cellular trajectories and molecular mechanisms of iPSC reprogramming. Curr Opin Genet Dev 52, 77–85.

6. Baluapuri, A., Hofstetter, J., Dudvarski Stankovic, N., Endres, T., Bhandare, P., Vos, S.M., Adhikari, B., Schwarz, J.D., Narain, A., Vogt, M., et al. (2019). MYC Recruits SPT5 to RNA Polymerase II to Promote Processive Transcription Elongation. Mol Cell 74, 674–687 e611.

7. Baluapuri, A., Wolf, E., and Eilers, M. (2020). Target gene-independent functions of MYC oncoproteins. Nat Rev Mol Cell Biol 21, 255–267.

8. Bartkova, J., Horejsi, Z., Koed, K., Kramer, A., Tort, F., Zieger, K., Guldberg, P., Sehested, M., Nesland, J.M., Lukas, C., et al. (2005). DNA damage response as a candidate anti-cancer barrier in early human tumorigenesis. Nature 434, 864–870.

9. Bertolini, M., Fenzl, K., Kats, I., Wruck, F., Tippmann, F., Schmitt, J., Auburger, J.J., Tans, S., Bukau, B., and Kramer, G. (2021). Interactions between nascent proteins translated by adjacent ribosomes drive homomer assembly. Science 371, 57–64.

10. Blackwood, E.M., Luscher, B., and Eisenman, R.N. (1992). Myc and Max associate in vivo. Genes Dev 6, 71–80.

11. Boija, A., Klein, I.A., Sabari, B.R., Dall’Agnese, A., Coffey, E.L., Zamudio, A.V., Li, C.H., Shrinivas, K., Manteiga, J.C., Hannett, N.M., et al. (2018). Transcription Factors Activate Genes through the Phase-Separation Capacity of Their Activation Domains. Cell 175, 1842–1855 e1816.

12. Brehme, M., Voisine, C., Rolland, T., Wachi, S., Soper, J.H., Zhu, Y., Orton, K., Villella, A., Garza, D., Vidal, M., et al. (2014). A chaperome subnetwork safeguards proteostasis in aging and neurodegenerative disease. Cell Rep 9, 1135–1150.

13. Brennan, C.M., Vaites, L.P., Wells, J.N., Santaguida, S., Paulo, J.A., Storchova, Z., Harper, J.W., Marsh, J.A., and Amon, A. (2019). Protein aggregation mediates stoichiometry of protein complexes in aneuploid cells. Genes Dev 33, 1031–1047.

14. Busch, S.J., and Sassone-Corsi, P. (1990). Dimers, leucine zippers and DNA-binding domains. Trends Genet 6, 36–40.

15. Carmel-Gross, I., Levy, E., Armon, L., Yaron, O., Waldman Ben-Asher, H., and Urbach, A. (2020). Human Pluripotent Stem Cell Fate Regulation by SMARCB1. Stem Cell Reports 15, 1037–1046.

16. Chen, J., Zhang, Z., Li, L., Chen, B.C., Revyakin, A., Hajj, B., Legant, W., Dahan, M., Lionnet, T., Betzig, E., et al. (2014). Single-molecule dynamics of enhanceosome assembly in embryonic stem cells. Cell 156, 1274–1285.

17. Chong, S., Graham, T.G.W., Dugast-Darzacq, C., Dailey, G.M., Darzacq, X., and Tjian, R. (2022). Tuning levels of low-complexity domain interactions to modulate endogenous oncogenic transcription. Mol Cell 82, 2084–2097 e2085.

18. Costa-Mattioli, M., and Walter, P. (2020). The integrated stress response: From mechanism to disease. Science 368.

19. Endres, T., Solvie, D., Heidelberger, J.B., Andrioletti, V., Baluapuri, A., Ade, C.P., Muhar, M., Eilers, U., Vos, S.M., Cramer, P., et al. (2021). Ubiquitylation of MYC couples transcription elongation with double-strand break repair at active promoters. Mol Cell 81, 830–844 e813.

20. Ferdous, A., Gonzalez, F., Sun, L., Kodadek, T., and Johnston, S.A. (2001). The 19S regulatory particle of the proteasome is required for efficient transcription elongation by RNA polymerase II. Mol Cell 7, 981–991.

21. Frottin, F., Schueder, F., Tiwary, S., Gupta, R., Korner, R., Schlichthaerle, T., Cox, J., Jungmann, R., Hartl, F.U., and Hipp, M.S. (2019). The nucleolus functions as a phase-separated protein quality control compartment. Science 365, 342–347.

22. Fu, A., Cohen-Kaplan, V., Avni, N., Livneh, I., and Ciechanover, A. (2021). p62-containing, proteolytically active nuclear condensates, increase the efficiency of the ubiquitin-proteasome system. Proc Natl Acad Sci U S A 118.

23. Greber, B.J., and Nogales, E. (2019). The Structures of Eukaryotic Transcription Pre-initiation Complexes and Their Functional Implications. Subcell Biochem 93, 143–192.

24. Guardavaccaro, D., Frescas, D., Dorrello, N.V., Peschiaroli, A., Multani, A.S., Cardozo, T., Lasorella, A., Iavarone, A., Chang, S., Hernando, E., et al. (2008). Control of chromosome stability by the beta-TrCP-REST-Mad2 axis. Nature 452, 365–369.

25. Guarnaccia, A.D., and Tansey, W.P. (2018). Moonlighting with WDR5: A Cellular Multitasker. J Clin Med 7.

26. Guo, X., Aviles, G., Liu, Y., Tian, R., Unger, B.A., Lin, Y.T., Wiita, A.P., Xu, K., Correia, M.A., and Kampmann, M. (2020). Mitochondrial stress is relayed to the cytosol by an OMA1-DELE1-HRI pathway. Nature 579, 427–432.

27. Harper, J.W., and Bennett, E.J. (2016). Proteome complexity and the forces that drive proteome imbalance. Nature 537, 328–338.

28. Harper, J.W., and Schulman, B.A. (2021). Cullin-RING Ubiquitin Ligase Regulatory Circuits: A Quarter Century Beyond the F-Box Hypothesis. Annu Rev Biochem 90, 403–429.

29. Heidelberger, J.B., Voigt, A., Borisova, M.E., Petrosino, G., Ruf, S., Wagner, S.A., and Beli, P. (2018). Proteomic profiling of VCP substrates links VCP to K6-linked ubiquitylation and c-Myc function. EMBO Rep 19.

30. Hipp, M.S., Park, S.H., and Hartl, F.U. (2014). Proteostasis impairment in protein-misfolding and -aggregation diseases. Trends Cell Biol 24, 506–514.

31. Hishida, T., Nozaki, Y., Nakachi, Y., Mizuno, Y., Okazaki, Y., Ema, M., Takahashi, S., Nishimoto, M., and Okuda, A. (2011). Indefinite self-renewal of ESCs through Myc/Max transcriptional complex-independent mechanisms. Cell stem cell 9, 37–49.

32. Hong, S., Choi, S., Kim, R., and Koh, J. (2020). Mechanisms of Macromolecular Interactions Mediated by Protein Intrinsic Disorder. Mol Cells 43, 899–908.

33. Hu, S., Peng, L., Xu, C., Wang, Z., Song, A., and Chen, F.X. (2021a). SPT5 stabilizes RNA polymerase II, orchestrates transcription cycles, and maintains the enhancer landscape. Mol Cell 81, 4425–4439 e4426.

34. Hu, X., Yu, J., Lin, Z., Feng, R., Wang, Z.W., and Chen, G. (2021b). The emerging role of WWP1 in cancer development and progression. Cell Death Discov 7, 163.

35. Huang, W., Pi, L., Liang, W., Xu, B., Wang, H., Cai, R., and Huang, H. (2006). The proteolytic function of the Arabidopsis 26S proteasome is required for specifying leaf adaxial identity. Plant Cell 18, 2479–2492.

36. Huber, A.L., Papp, S.J., Chan, A.B., Henriksson, E., Jordan, S.D., Kriebs, A., Nguyen, M., Wallace, M., Li, Z., Metallo, C.M., et al. (2016). CRY2 and FBXL3 Cooperatively Degrade c-MYC. Mol Cell 64, 774–789.

37. Huttlin, E.L., Bruckner, R.J., Navarrete-Perea, J., Cannon, J.R., Baltier, K., Gebreab, F., Gygi, M.P., Thornock, A., Zarraga, G., Tam, S., et al. (2021). Dual proteome-scale networks reveal cell-specific remodeling of the human interactome. Cell 184, 3022–3040 e3028.

38. Jackson, D.A., Hassan, A.B., Errington, R.J., and Cook, P.R. (1993). Visualization of focal sites of transcription within human nuclei. EMBO J 12, 1059–1065.

39. Jimenez Martin, O., Schlosser, A., Furtwangler, R., Wegert, J., and Gessler, M. (2021). MYCN and MAX alterations in Wilms tumor and identification of novel N-MYC interaction partners as biomarker candidates. Cancer Cell Int 21, 555.

40. Jungblut, A., Hopfner, K.P., and Eustermann, S. (2020). Megadalton chromatin remodelers: common principles for versatile functions. Curr Opin Struct Biol 64, 134–144.

41. Juszkiewicz, S., and Hegde, R.S. (2018). Quality Control of Orphaned Proteins. Mol Cell 71, 443–457.

42. Kaushik, S., and Cuervo, A.M. (2015). Proteostasis and aging. Nat Med 21, 1406–1415.

43. Kim, J.G., Shin, H.C., Seo, T., Nawale, L., Han, G., Kim, B.Y., Kim, S.J., and Cha-Molstad, H. (2021). Signaling Pathways Regulated by UBR Box-Containing E3 Ligases. Int J Mol Sci 22.

44. King, B., Boccalatte, F., Moran-Crusio, K., Wolf, E., Wang, J., Kayembe, C., Lazaris, C., Yu, X., Aranda-Orgilles, B., Lasorella, A., et al. (2016). The ubiquitin ligase Huwe1 regulates the maintenance and lymphoid commitment of hematopoietic stem cells. Nat Immunol 17, 1312–1321.

45. Kinyamu, H.K., and Archer, T.K. (2007). Proteasome activity modulates chromatin modifications and RNA polymerase II phosphorylation to enhance glucocorticoid receptor-mediated transcription. Mol Cell Biol 27, 4891–4904.

46. Kolla, S., Ye, M., Mark, K.G., and Rape, M. (2022). Assembly and function of branched ubiquitin chains. Trends Biochem Sci 47, 759–771.

47. Koren, I., Timms, R.T., Kula, T., Xu, Q., Li, M.Z., and Elledge, S.J. (2018). The Eukaryotic Proteome Is Shaped by E3 Ubiquitin Ligases Targeting C-Terminal Degrons. Cell 173, 1622–1635 e1614.

48. Li, J., Cai, Z., Vaites, L.P., Shen, N., Mitchell, D.C., Huttlin, E.L., Paulo, J.A., Harry, B.L., and Gygi, S.P. (2021). Proteome-wide mapping of short-lived proteins in human cells. Mol Cell.

49. Liao, L., Song, M., Li, X., Tang, L., Zhang, T., Zhang, L., Pan, Y., Chouchane, L., and Ma, X. (2017). E3 Ubiquitin Ligase UBR5 Drives the Growth and Metastasis of Triple-Negative Breast Cancer. Cancer Res 77, 2090–2101.

50. Littler, S., Sloss, O., Geary, B., Pierce, A., Whetton, A.D., and Taylor, S.S. (2019). Oncogenic MYC amplifies mitotic perturbations. Open Biol 9, 190136.

51. Liu, X., Reitsma, J.M., Mamrosh, J.L., Zhang, Y., Straube, R., and Deshaies, R.J. (2018). Cand1-Mediated Adaptive Exchange Mechanism Enables Variation in F-Box Protein Expression. Mol Cell 69, 773–786 e776.

52. Lourenco, C., Resetca, D., Redel, C., Lin, P., MacDonald, A.S., Ciaccio, R., Kenney, T.M.G., Wei, Y., Andrews, D.W., Sunnerhagen, M., et al. (2021). MYC protein interactors in gene transcription and cancer. Nat Rev Cancer 21, 579–591.

53. Lu, H., Yu, D., Hansen, A.S., Ganguly, S., Liu, R., Heckert, A., Darzacq, X., and Zhou, Q. (2018). Phase-separation mechanism for C-terminal hyperphosphorylation of RNA polymerase II. Nature 558, 318–323.

54. Magits, W., and Sablina, A.A. (2022). The regulation of the protein interaction network by monoubiquitination. Curr Opin Struct Biol 73, 102333.

55. Manford, A.G., Mena, E.L., Shih, K.Y., Gee, C.L., McMinimy, R., Martinez-Gonzalez, B., Lew, B.G., Zoltek, M., Rodriguez-Perez, F., Woldesenbet, M., et al. (2021). Structural basis and regulation of the reductive stress response. Cell in revision.

56. Manford, A.G., Rodriguez-Perez, F., Shih, K.Y., Shi, Z., Berdan, C.A., Choe, M., Titov, D.V., Nomura, D.K., and Rape, M. (2020). A Cellular Mechanism to Detect and Alleviate Reductive Stress. Cell 183, 46–61 e21.

57. Meacham, G.C., Patterson, C., Zhang, W., Younger, J.M., and Cyr, D.M. (2001). The Hsc70 co-chaperone CHIP targets immature CFTR for proteasomal degradation. Nat Cell Biol 3, 100–105.

58. Meissner, B., Kridel, R., Lim, R.S., Rogic, S., Tse, K., Scott, D.W., Moore, R., Mungall, A.J., Marra, M.A., Connors, J.M., et al. (2013). The E3 ubiquitin ligase UBR5 is recurrently mutated in mantle cell lymphoma. Blood 121, 3161–3164.

59. Mena, E.L., Jevtic, P., Greber, B.J., Gee, C.L., Lew, B.G., Akopian, D., Nogales, E., Kuriyan, J., and Rape, M. (2020). Structural basis for dimerization quality control. Nature 586, 452–456.

60. Mena, E.L., Kjolby, R.A.S., Saxton, R.A., Werner, A., Lew, B.G., Boyle, J.M., Harland, R., and Rape, M. (2018). Dimerization quality control ensures neuronal development and survival. Science 362.

61. Meyer, H.J., and Rape, M. (2014). Enhanced protein degradation by branched ubiquitin chains. Cell 157, 910–921.

62. Oh, E., Mark, K.G., Mocciaro, A., Watson, E.R., Prabu, J.R., Cha, D.D., Kampmann, M., Gamarra, N., Zhou, C.Y., and Rape, M. (2020). Gene expression and cell identity controlled by anaphase-promoting complex. Nature 579, 136–140.

63. Ohtake, F., Tsuchiya, H., Saeki, Y., and Tanaka, K. (2018). K63 ubiquitylation triggers proteasomal degradation by seeding branched ubiquitin chains. Proc Natl Acad Sci U S A.

64. Osborne, C.S., Chakalova, L., Mitchell, J.A., Horton, A., Wood, A.L., Bolland, D.J., Corcoran, A.E., and Fraser, P. (2007). Myc dynamically and preferentially relocates to a transcription factory occupied by Igh. PLoS Biol 5, e192.

65. Padovani, C., Jevtic, P., and Rape, M. (2022). Quality control of protein complex composition. Mol Cell 82, 1439–1450.

66. Papantonis, A., and Cook, P.R. (2013). Transcription factories: genome organization and gene regulation. Chem Rev 113, 8683–8705.

67. Patel, A., Lee, H.O., Jawerth, L., Maharana, S., Jahnel, M., Hein, M.Y., Stoynov, S., Mahamid, J., Saha, S., Franzmann, T.M., et al. (2015). A Liquid-to-Solid Phase Transition of the ALS Protein FUS Accelerated by Disease Mutation. Cell 162, 1066–1077.

68. Pierce, N.W., Kleiger, G., Shan, S.O., and Deshaies, R.J. (2009). Detection of sequential polyubiquitylation on a millisecond timescale. Nature 462, 615–619.

69. Qiao, X., Liu, Y., Prada, M.L., Mohan, A.K., Gupta, A., Jaiswal, A., Sharma, M., Merisaari, J., Haikala, H.M., Talvinen, K., et al. (2020). UBR5 Is Coamplified with MYC in Breast Tumors and Encodes an Ubiquitin Ligase That Limits MYC-Dependent Apoptosis. Cancer Res 80, 1414–1427.

70. Reavie, L., Della Gatta, G., Crusio, K., Aranda-Orgilles, B., Buckley, S.M., Thompson, B., Lee, E., Gao, J., Bredemeyer, A.L., Helmink, B.A., et al. (2010). Regulation of hematopoietic stem cell differentiation by a single ubiquitin ligase-substrate complex. Nat Immunol 11, 207–215.

71. Rippe, K., and Papantonis, A. (2022). Functional organization of RNA polymerase II in nuclear subcompartments. Curr Opin Cell Biol 74, 88–96.

72. Rodrigo-Brenni, M.C., Gutierrez, E., and Hegde, R.S. (2014). Cytosolic quality control of mislocalized proteins requires RNF126 recruitment to Bag6. Mol Cell 55, 227–237.

73. Salghetti, S.E., Caudy, A.A., Chenoweth, J.G., and Tansey, W.P. (2001). Regulation of transcriptional activation domain function by ubiquitin. Science 293, 1651–1653.

74. Salghetti, S.E., Muratani, M., Wijnen, H., Futcher, B., and Tansey, W.P. (2000). Functional overlap of sequences that activate transcription and signal ubiquitin-mediated proteolysis. Proc Natl Acad Sci U S A 97, 3118–3123.

75. Saunders, D.N., Hird, S.L., Withington, S.L., Dunwoodie, S.L., Henderson, M.J., Biben, C., Sutherland, R.L., Ormandy, C.J., and Watts, C.K. (2004). Edd, the murine hyperplastic disc gene, is essential for yolk sac vascularization and chorioallantoic fusion. Mol Cell Biol 24, 7225–7234.

76. Scaglione, K.M., Zavodszky, E., Todi, S.V., Patury, S., Xu, P., Rodriguez-Lebron, E., Fischer, S., Konen, J., Djarmati, A., Peng, J., et al. (2011). Ube2w and Ataxin-3 Coordinately Regulate the Ubiquitin Ligase CHIP. Mol Cell 43, 599–612.

77. Schukur, L., Zimmermann, T., Niewoehner, O., Kerr, G., Gleim, S., Bauer-Probst, B., Knapp, B., Galli, G.G., Liang, X., Mendiola, A., et al. (2020). Identification of the HECT E3 ligase UBR5 as a regulator of MYC degradation using a CRISPR/Cas9 screen. Sci Rep 10, 20044.

78. Siegel, J.J., and Amon, A. (2012). New insights into the troubles of aneuploidy. Annu Rev Cell Dev Biol 28, 189–214.

79. Sievers, Q.L., Petzold, G., Bunker, R.D., Renneville, A., Slabicki, M., Liddicoat, B.J., Abdulrahman, W., Mikkelsen, T., Ebert, B.L., and Thoma, N.H. (2018). Defining the human C2H2 zinc finger degrome targeted by thalidomide analogs through CRBN. Science 362.

80. Snead, W.T., and Gladfelter, A.S. (2019). The Control Centers of Biomolecular Phase Separation: How Membrane Surfaces, PTMs, and Active Processes Regulate Condensation. Mol Cell 76, 295–305.

81. Song, M., Yeku, O.O., Rafiq, S., Purdon, T., Dong, X., Zhu, L., Zhang, T., Wang, H., Yu, Z., Mai, J., et al. (2020). Tumor derived UBR5 promotes ovarian cancer growth and metastasis through inducing immunosuppressive macrophages. Nat Commun 11, 6298.

82. Sotillo, R., Hernando, E., Diaz-Rodriguez, E., Teruya-Feldstein, J., Cordon-Cardo, C., Lowe, S.W., and Benezra, R. (2007). Mad2 overexpression promotes aneuploidy and tumorigenesis in mice. Cancer Cell 11, 9–23.

83. Sotillo, R., Schvartzman, J.M., Socci, N.D., and Benezra, R. (2010). Mad2-induced chromosome instability leads to lung tumour relapse after oncogene withdrawal. Nature 464, 436–440.

84. Tan, X., Calderon-Villalobos, L.I., Sharon, M., Zheng, C., Robinson, C.V., Estelle, M., and Zheng, N. (2007). Mechanism of auxin perception by the TIR1 ubiquitin ligase. Nature 446, 640–645.

85. Tastemel, M., Gogate, A.A., Malladi, V.S., Nguyen, K., Mitchell, C., Banaszynski, L.A., and Bai, X. (2017). Transcription pausing regulates mouse embryonic stem cell differentiation. Stem Cell Res 25, 250–255.

86. Thomas, L.R., Wang, Q., Grieb, B.C., Phan, J., Foshage, A.M., Sun, Q., Olejniczak, E.T., Clark, T., Dey, S., Lorey, S., et al. (2015). Interaction with WDR5 promotes target gene recognition and tumorigenesis by MYC. Mol Cell 58, 440–452.

87. Torres, E.M., Sokolsky, T., Tucker, C.M., Chan, L.Y., Boselli, M., Dunham, M.J., and Amon, A. (2007). Effects of aneuploidy on cellular physiology and cell division in haploid yeast. Science 317, 916–924.

88. Toyama, B.H., Savas, J.N., Park, S.K., Harris, M.S., Ingolia, N.T., Yates, J.R, 3rd., and Hetzer, M.W. (2013). Identification of long-lived proteins reveals exceptional stability of essential cellular structures. Cell 154, 971-982.

89. Trojanowski, J., Frank, L., Rademacher, A., Mucke, N., Grigaitis, P., and Rippe, K. (2022). Transcription activation is enhanced by multivalent interactions independent of phase separation. Mol Cell 82, 1878–1893 e1810.

90. Vilchez, D., Simic, M.S., and Dillin, A. (2014). Proteostasis and aging of stem cells. Trends Cell Biol 24, 161–170.

91. Wagh, K., Garcia, D.A., and Upadhyaya, A. (2021). Phase separation in transcription factor dynamics and chromatin organization. Curr Opin Struct Biol 71, 148–155.

92. Wang, L., Du, Y., Ward, J.M., Shimbo, T., Lackford, B., Zheng, X., Miao, Y.L., Zhou, B., Han, L., Fargo, D.C., et al. (2014). INO80 facilitates pluripotency gene activation in embryonic stem cell self-renewal, reprogramming, and blastocyst development. Cell stem cell 14, 575–591.

93. Wang, X., Arceci, A., Bird, K., Mills, C.A., Choudhury, R., Kernan, J.L., Zhou, C., Bae-Jump, V., Bowers, A., and Emanuele, M.J. (2017). VprBP/DCAF1 Regulates the Degradation and Nonproteolytic Activation of the Cell Cycle Transcription Factor FoxM1. Mol Cell Biol 37.

94. Wei, M.T., Chang, Y.C., Shimobayashi, S.F., Shin, Y., Strom, A.R., and Brangwynne, C.P. (2020). Nucleated transcriptional condensates amplify gene expression. Nat Cell Biol 22, 1187–1196.

95. Weissmiller, A.M., Wang, J., Lorey, S.L., Howard, G.C., Martinez, E., Liu, Q., and Tansey, W.P. (2019). Inhibition of MYC by the SMARCB1 tumor suppressor. Nat Commun 10, 2014.

96. Welcker, M., and Clurman, B.E. (2008). FBW7 ubiquitin ligase: a tumour suppressor at the crossroads of cell division, growth and differentiation. Nat Rev Cancer 8, 83–93.

97. Welcker, M., Orian, A., Jin, J., Grim, J.E., Harper, J.W., Eisenman, R.N., and Clurman, B.E. (2004). The Fbw7 tumor suppressor regulates glycogen synthase kinase 3 phosphorylation-dependent c-Myc protein degradation. Proc Natl Acad Sci U S A 101, 9085–9090.

98. Welcker, M., Wang, B., Rusnac, D.V., Hussaini, Y., Swanger, J., Zheng, N., and Clurman, B.E. (2022). Two diphosphorylated degrons control c-Myc degradation by the Fbw7 tumor suppressor. Sci Adv 8, eabl7872.

99. Wenzel, D.M., Lissounov, A., Brzovic, P.S., and Klevit, R.E. (2011). UBCH7 reactivity profile reveals parkin and HHARI to be RING/HECT hybrids. Nature 474, 105–108.

100. Werner, A., Iwasaki, S., McGourty, C.A., Medina-Ruiz, S., Teerikorpi, N., Fedrigo, I., Ingolia, N.T., and Rape, M. (2015). Cell-fate determination by ubiquitin-dependent regulation of translation. Nature 525, 523–527.

101. Wickliffe, K.E., Lorenz, S., Wemmer, D.E., Kuriyan, J., and Rape, M. (2011). The mechanism of linkage-specific ubiquitin chain elongation by a single-subunit e2. Cell 144, 769–781.

102. Williamson, A., Wickliffe, K.E., Mellone, B.G., Song, L., Karpen, G.H., and Rape, M. (2009). Identification of a physiological E2 module for the human anaphase-promoting complex. Proc Natl Acad Sci U S A 106, 18213–18218.

103. Wu, R.C., Feng, Q., Lonard, D.M., and O’Malley, B.W. (2007). SRC-3 coactivator functional lifetime is regulated by a phospho-dependent ubiquitin time clock. Cell 129, 1125–1140.

104. Yao, R., Zhang, M., Zhou, J., Liu, L., Zhang, Y., Gao, J., and Xu, K. (2022). Novel dual-targeting c-Myc inhibitor D347-2761 represses myeloma growth via blocking c-Myc/Max heterodimerization and disturbing its stability. Cell Commun Signal 20, 73.

105. Yau, R., and Rape, M. (2016). The increasing complexity of the ubiquitin code. Nat Cell Biol 18, 579–586.

106. Yau, R.G., Doerner, K., Castellanos, E.R., Haakonsen, D.L., Werner, A., Wang, N., Yang, X.W., Martinez-Martin, N., Matsumoto, M.L., Dixit, V.M., et al. (2017). Assembly and Function of Heterotypic Ubiquitin Chains in Cell-Cycle and Protein Quality Control. Cell 171, 918–933 e920.

107. Zavodszky, E., Peak-Chew, S.Y., Juszkiewicz, S., Narvaez, A.J., and Hegde, R.S. (2021). Identification of a quality-control factor that monitors failures during proteasome assembly. Science 373, 998–1004.

108. Zhou, B., Wang, L., Zhang, S., Bennett, B.D., He, F., Zhang, Y., Xiong, C., Han, L., Diao, L., Li, P., et al. (2016). INO80 governs superenhancer-mediated oncogenic transcription and tumor growth in melanoma. Genes Dev 30, 1440–1453.

